# The receptor kinase NILR1 confers nematode resistance through developmental rather than canonical immune signaling

**DOI:** 10.64898/2026.07.24.740501

**Authors:** Maximilian F. Euler, Seema Aslam, Stefan Neumann, Florian M.W. Grundler

## Abstract

- NILR1 contributes to quantitative resistance against *Heterodera schachtii* and signals through a brassinosteroid-type kinase cascade, but whether its contribution engages canonical immune signaling or follows a distinct, developmentally biased logic has remained unclear.
- We profiled early transcriptional responses of wild-type and *nilr1* roots to *H. schachtii* by RNA-seq at 30 min and 3 h post inoculation and analyzed them using a genotype × treatment interaction model, complemented by gene set enrichment and integration with cell-type-resolved, brassinosteroid-responsive regulatory networks.
- We found that canonical PTI marker transcription and flg22-induced ROS production were preserved or even elevated in *nilr1*, indicating that the tested PTI outputs are largely maintained despite increased susceptibility. Instead, genes whose nematode-induced responses showed NILR1-sensitive interaction effects were enriched for a HAT7/GTL1 centred cortex developmental module, and genetic disruption of GTL1/DF1 moderately altered *H. schachtii* parasitism, consistent with a role for this module in shaping host permissiveness.
- Our data support a model in which NILR1 contributes to nematode resistance primarily via developmental, brassinosteroid-type signaling rather than via the canonical PTI outputs assayed here, consistent with a co-option of BRI1 clade (LRR-RLK-Xb) architecture — a receptor family classically associated with development — into a pathway that modulates susceptibility to a plant-parasitic nematode.

## Introduction

Plant-parasitic nematodes are among the most damaging soilborne pathogens of major crops (Savary et al., 2019), yet the receptor-level signaling events that govern early host responses to root invasion remain poorly understood. Plants perceive and respond to pathogens through plasma membrane–localized pattern-recognition receptors (PRRs) which detect conserved molecular signatures and activate pattern-triggered immunity (PTI; Jones and Dangl, 2006; Ngou et al., 2026). PTI signaling has been extensively characterized for microbial elicitors such as bacterial flagellin (Felix et al., 1999; Chinchilla et al., 2006) and fungal chitin (Miya et al., 2007). More recently, a nematode ligand–receptor pair was described in which a trehalase-derived molecular pattern is recognized by *LecRK-V.5/V.6* (Iino et al., 2025). During early infection, however, migratory cyst nematodes traverse cortical root tissues before initiating feeding sites near the vascular cylinder (Wyss and Zunke, 1986; Siddique and Grundler, 2018), raising the possibility that the transcriptional programs governing cortical cell growth, identity, and remodeling are themselves determinants of early host susceptibility – a view supported by the recent finding that the HD-ZIP transcription factor *AtATHB2* and its rice ortholog *OsHOX28* are functionally required for gall formation by root-knot nematodes (Saura-Sanchez et al., 2025, *bioRxiv*). Nevertheless, the receptors and transcriptional programs that mediate early root responses to nematode invasion remain largely undefined.

In *Arabidopsis thaliana*, the leucine-rich repeat receptor-like kinase *NEMATODE-INDUCED LRR-RLK 1* (*NILR1*) is induced during infection by the cyst nematode *Heterodera schachtii*, and loss-of-function mutants show increased susceptibility (Mendy et al., 2017). NILR1 contributes to PTI-associated immune responses and functions with the co-receptor BAK1 (Mendy et al., 2017). Recently, the conserved nematode pheromone ascaroside-18 (Ascr18) was identified as a NILR1-perceived molecular pattern that triggers immune signaling and resistance (Huang et al., 2023), although Ascr18-triggered defense can also proceed through NILR1-independent repression of auxin signaling (Letia et al., 2025) or NILR1-dependent priming of defense gene loci for enhanced activation upon pathogen challenge (Manohar et al., 2026). In parallel, NILR1 was characterized as a growth-associated receptor kinase within the BRI1 family that engages a conserved brassinosteroid-type intracellular signaling cascade (Z. Wu et al., 2017; Zheng et al., 2022; Ali et al., 2025). This subfamily is classically associated with development rather than immunity, in contrast to the LRR-RLK clades that activate MAPK-driven defense transcription (Jones et al., 2024; Ngou et al., 2026). Whether NILR1 activates canonical immune transcription despite this developmental architecture, or instead shapes infection-induced gene expression through a different regulatory logic, remains unresolved.

Existing transcriptomic studies of cyst nematode infection have predominantly sampled post-invasion time points, when migrating juveniles have already entered the root tissue and feeding-site reprogramming begins to dominate the transcriptional landscape (Mendy et al., 2017; Siddique et al., 2022; Pijnacker et al., 2026; Saura-Sanchez et al., 2025). Resolving receptor-level events therefore requires sampling that captures both the earliest perception phase, when infective juveniles approach and contact the root surface, and the immediate downstream phase, in which receptor activation is translated into transcriptional output.

Here, we characterize NILR1-associated differences in transcriptional reprogramming during early *H. schachtii* infection using time-resolved RNA sequencing at 30 min and 3 h post inoculation, combined with a genotype × treatment interaction framework. In this framework, we find that nematode-induced activation of canonical PTI marker genes is preserved or enhanced in *nilr1* roots, indicating that the canonical PTI transcriptional program assessed here does not require NILR1. By contrast, genes with significant or shrinkage-prioritised genotype × treatment interaction effects are enriched for coordinated defense-and remodeling-associated programs that overlap with the HAT7/GTL1 cortex regulatory network described by Nolan et al., 2023. Consistent with this network-level signal, genetic disruption of the GTL1/DF1 transcription factor module led to moderate, genotype-specific changes in *H. schachtii* parasitism in our assays, supporting a contribution of this developmental module to host permissiveness. Together, these findings support a model in which NILR1 functions as a developmentally wired receptor whose activation tunes a pre-existing cortex regulatory module, and they illustrate that increased resistance to a plant-parasitic nematode can, at least in this system, coincide with largely intact canonical immune outputs.

## Materials and Methods

### Plant material and growth conditions

Seeds of *Arabidopsis thaliana* (L.) Heynh. ecotype Columbia-0 (Col-0), *nilr1* (AT1G74360; T-DNA insertion line SAIL 865 E07, corresponds to *nilr1-1* in Huang et al., 2023 and *grace-1* in Z. Wu et al., 2017), *gtl1-1* (WiscDsLox413-416C9), and the *gtl1-1 df1-1* double mutant (Shibata et al., 2018) were surface-sterilized by immersion in 96% (v/v) ethanol for 1 min, followed by sodium hypochlorite treatment for 3 min (1.3% NaOCl for Col-0 and 0.6% NaOCl for the mutants). Seeds were subsequently rinsed three times with sterile distilled water for 2 min each and dried on sterile filter paper.

### Infection assays

Sterilized seeds were germinated on Knop medium under long days (16 h light / 8 h dark, 23 ± 1*^◦^*C). Cysts of *Heterodera schachtii* ’Bonn’, maintained on sterile mustard (*Sinapis alba cv. Albatros*), were hatched in a Baermann funnel with 3 mM ZnCl_2_. Twelve-day-old seedlings (≥ 20 per genotype) were inoculated with ca. 80 freshly hatched J2s. Males and females were counted at 12 d post-inoculation (dpi); syncytia and female size were measured at 14 dpi (Leica M165C, LAS v. 4.3). *n* ≥ 3 replicates per assay.

For Col-0 vs *nilr1* (Fig. 1; Supplementary Table S13), analyses used Python (statsmodels, SciPy). Counts (female, male, total) were analyzed with negative-binomial GLMs (Count ∼ Genotype + Replicate; replicate as fixed block; genotype effects are reported as incidence rate ratios with 95% confidence intervals), with consistency checked by the genotype × replicate interaction and paired *t*-tests (*n* = 3). Female and syncytium sizes were fitted analogously on log scale (geometric-mean ratios). *P <* 0.05 was considered significant.

**Figure 1:**
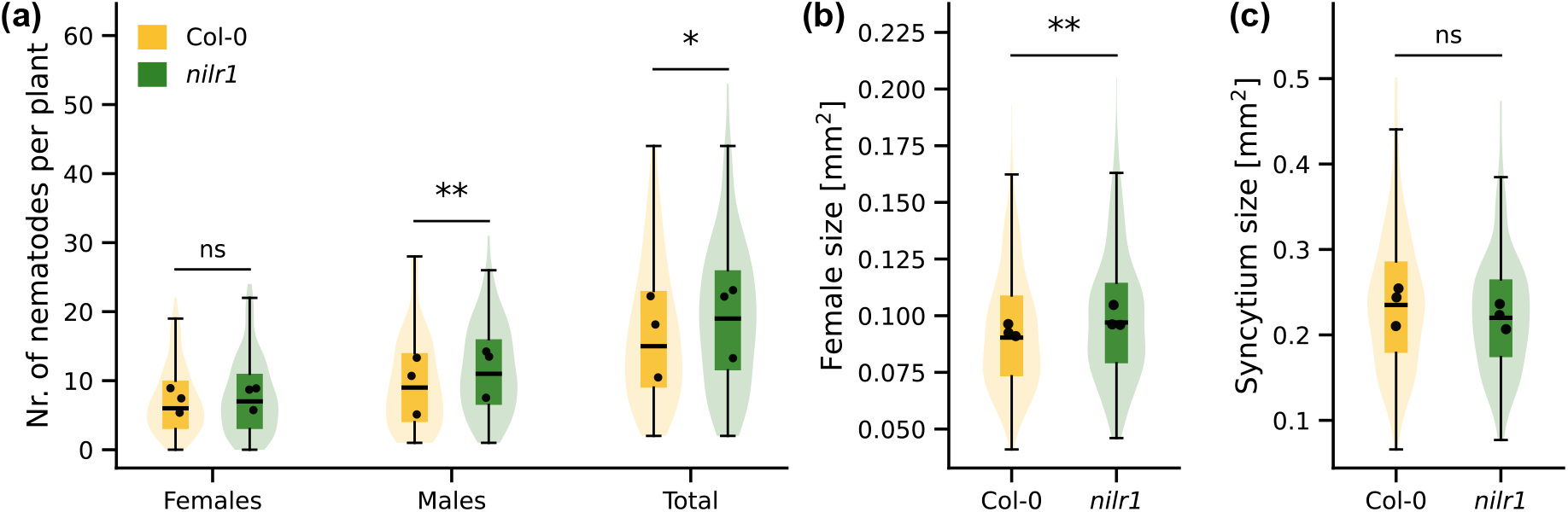
Phenotype of Col-0 and *nilr1* under infection with *H. schachtii*. **(a)** Number of female, male, and total *H. schachtii* per plant at 12 days post inoculation (dpi). Violin plots show the distribution of plant-level counts, with overlaid boxplots indicating the median and interquartile range; black dots represent biological replicate means (n = 3 independent biological replicates, ≥ 36 plants per genotype across replicates). Statistical significance was assessed using negative-binomial generalized linear models with genotype as a fixed effect and biological replicate as a fixed blocking factor (Count ∼ Genotype + Replicate), accounting for between-experiment variation in baseline infection. Female **(b)** and syncytium sizes **(c)** in Col-0 and *nilr1* roots, measured at 14 dpi. Statistical significance was assessed using linear models on log-transformed measurements with the same fixed-block structure (log(Size) ∼ Genotype + Replicate). Significance levels are indicated as *^∗^P <* 0.05, *^∗∗^P <* 0.01, *^∗∗∗^P <* 0.001, and ns (*P* ≥ 0.05).

For the multi-genotype comparison (Fig. 5; Supplementary Table S14), counts were fitted in R by a negative-binomial GLMM (glmmTMB; Genotype × Sex fixed, plant-in-replicate random intercepts; Type II Wald *χ*^2^ tests). Pairwise comparisons were performed within each sex using estimated marginal means with Tukey adjustment (emmeans). Total counts were analyzed with an analogous Genotype-only model. Female and syncytium sizes (log scale) used linear mixed models (lme4, lmerTest; Tukey EMMs, Satterthwaite df). Compact letter displays at *α* = 0.05 were generated with S^̌^ idák adjustment (emmeans/multcomp).

### Plant growth and inoculation for RNA-seq

A balanced factorial RNA-seq experiment was performed with two genotypes (Col-0 and *nilr1* loss-of-function mutant), two treatments (mock and *H. schachtii* inoculation), and two time points (30 min and 3 h after inoculation). Plants were germinated in 6-well plates with 800 µl half-strength liquid MS medium (pH 6.5). After 7 days, three seedlings were transferred to each well of new 6-well plates filled with 1 ml half-strength liquid MS. For treatments, approximately 7,000 freshly hatched *H. schachtii* J2s were added per well to the 12-day-old plants. Wells receiving an equal volume of water served as mock controls. Roots were collected after 30 min and 3 h of incubation, gently blotted dry on sterile filter paper, excised with a scalpel, and immediately snap-frozen in liquid nitrogen. Total RNA was extracted from frozen root tissues using the Quick-RNA Plant Miniprep Kit (R2024, Zymo Research) following the manufacturer’s instructions. Residual genomic DNA was removed with the TURBO DNA-free Kit (Invitrogen) according to the supplied protocol. RNA quantity was assessed spectrophotometrically using a Nanodrop 2000c (Thermo Scientific). RNA integrity was evaluated by Novogene prior to library preparation.

The 10 h timepoint shown in Figures 2e and 5d was generated in a separate experiment using the same setup (wild-type only) and RNA-seq pipeline, and is shown solely as temporal context and does not contribute to the genotype × treatment interaction analysis.

**Figure 2:**
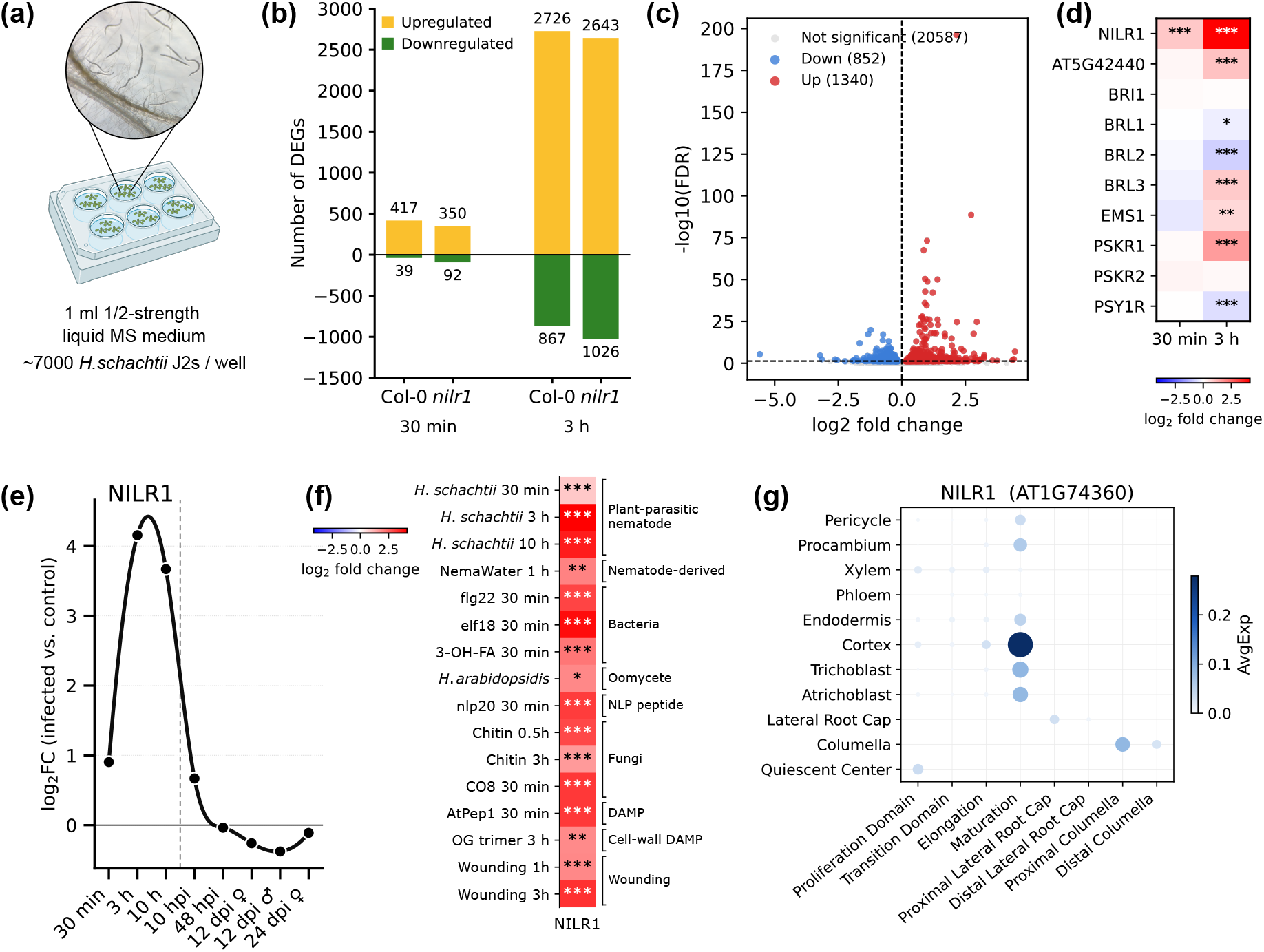
Experimental setup and early NILR1-dependent transcriptional responses to cyst nematode infection. **(a)** Wild-type (Col-0) and *nilr1* mutant *A. thaliana* seedlings were grown in liquid half-strength MS medium and inoculated with approximately 7,000 *H. schachtii* J2s or mock-treated. Roots were harvested after 30 min and 3 h for RNA-seq analysis. **(b)** Numbers of differentially expressed genes (DEGs) in Col-0 and *nilr1* roots at 30 min and 3 h after nematode inoculation relative to mock controls, separated into up- and downregulated genes. **(c)** Baseline expression (*nilr1* vs WT) showing 1340 de-repressed and 852 repressed genes in *nilr1* mutants. Adjusted *P <* 0.05, *n* = 8 per genotype. **(d)** Heatmap of *H. schachtii* –induced log2 fold changes (infected vs. mock) for *Arabidopsis* BRI1-clade (LRR-RLK-Xb) receptor kinases in wild-type roots at 30 min and 3 h post inoculation; asterisks denote DESeq2 adjusted *P* values (* *P <* 0.05, ** *P <* 0.01, *** *P <* 0.001). **(e)** Expression trajectory of the NILR1 receptor across early infection (30 min and 3 h from this study; 10 h from a separate experiment with identical setup) and the post-invasion lifecycle (Siddique et al., 2022; 10 hours post infection, hpi – 24 days post infection, dpi) in Col-0 wild-type roots. Values are average log2 fold changes of infected over mock-treated roots. **(f)** NILR1 is significantly upregulated as part of the core *A. thaliana* immune response (Data from *H. schachtii:* this study; NemaWater: Mendy et al., 2017; flg22, AtPep1: Safaeizadeh et al., 2024; *H. arabidopsidis*: Ramirez et al., 2026; Chitin: Stringlis et al., 2018; OG trimers: Davidsson et al., 2017; Wounding: Rymen et al., 2019; elf18, 3-OH-FA, nlp20, CO8: Bjornson et al., 2021). **(g)** Cell-type-resolved expression of *NILR1* across root cell types, based on published single-cell transcriptomic data (Shahan et al., 2022). *NILR1* shows highest expression in cortex cells.

### RNA-seq library preparation, sequencing, and read processing

Library preparation and sequencing were performed by Novogene (Munich, Germany). Messenger RNA (mRNA) was purified from total RNA by poly(A) enrichment using poly-T oligo-attached magnetic beads. The purified mRNA was fragmented and converted into first-strand cDNA using random hexamer primers, followed by second-strand synthesis, end repair, 3’ adenylation, adapter ligation, size selection, PCR amplification, and purification. Libraries were sequenced on an Illumina NovaSeq X Plus platform to generate 150 bp paired-end reads, yielding approximately 50 million read pairs per library. Each of the eight genotype × treatment × time combinations was represented by four independent biological replicates, except *nilr1* at 30 minutes with nematode inoculation, for which one replicate was excluded due to a low RNA integrity number (RIN), resulting in a total of 31 RNA-seq libraries. Raw reads were filtered by Novogene to remove adapter sequences, reads with more than 10% ambiguous bases, and reads with more than 50% low-quality bases (Phred score *<* 5). Clean reads were aligned to the *A. thaliana* TAIR10.1 reference genome (NCBI GCF 000001735.4) using HISAT2 v2.2.1 (Kim et al., 2019) with default parameters. Gene-level read counts were obtained with featureCounts v2.0.6 (Liao et al., 2014) using the corresponding NCBI gene annotation. Principal component analysis (Fig. S1) was performed by Novogene on FPKM values across all genes. Disruption of the NILR1 locus in the *nilr1* line was confirmed by RNA-seq coverage analysis (Supplementary Fig. S2).

### Differential expression and genotype **×** treatment interaction analysis

Differential expression analysis was performed using DESeq2 (Love et al., 2014) with a genotype × treatment interaction model (∼genotype + treatment + genotype:treatment), setting wild-type and control as reference levels; genes with fewer than ten total counts across all samples were excluded prior to testing. Because interaction terms are estimated with lower power than main effects, we prioritised candidates using the *apeglm* shrinkage estimator (Zhu et al., 2019, Fig. S3)—which collapses poorly supported effects toward zero while preserving tightly estimated ones—rather than a formal FDR threshold. Genes with shrunken log_2_ FC ≤ −0.2 were defined as NILR1-dependent candidates and those with shrunken log_2_ FC ≥ 0.2 as the complementary set of enhanced or derepressed responses. A negative interaction coefficient reflects an attenuated *H. schachtii* response in *nilr1* relative to wild type, encompassing absent induction (wild type induced, *nilr1* unchanged), destabilization (wild type stable, *nilr1* repressed), or reduced amplitude (both respond, *nilr1* weaker). This operational threshold identifies well-supported effects but does not constitute a calibrated false discovery procedure; full interaction statistics—unshrunken and shrunken log_2_ fold changes, Wald statistics, and adjusted *p*-values—for all tested genes are provided in Supplementary Tables S5 (30 min) and S6 (3 h).

### Generation of the pooled baseline dataset

The constitutive *nilr1*-versus-wild-type transcriptome was defined from the 16 control (mock-inoculated) libraries of the infection time course — four biological replicates of each genotype at each of two time points (30 min and 3 h), i.e. 8 wild-type and 8 *nilr1* samples. Raw, unnormalized gene-level counts (genes quantified in all libraries and with a summed count ≥ 10 retained) were analyzed in DESeq2. The two time points were treated as additional replication of the resting state and pooled in a single model with time point as a blocking factor (design ̃time + genotype; wild type and 30 min as reference levels), giving *n* = 8 per genotype under a common dispersion estimate. The constitutive genotype effect (*nilr1* vs wild type) was assessed by the Wald test with Benjamini–Hochberg FDR correction (significant at adjusted *P <* 0.05), and log_2_ fold changes were apeglm-shrunken for ranking and visualization. To verify that this pooled effect was stable rather than time-point-specific, a companion model including the genotype × time interaction (̃ time + genotype + time:genotype) was compared to the reduced model by a likelihood-ratio test, and genes with a significant interaction (adjusted *P <* 0.05) were flagged but retained.

### Gene set enrichment analysis

Rank-based gene set enrichment analysis (GSEA) was performed using the clusterProfiler package (v4.14.6, T. Wu et al., 2021) with the adaptive multilevel permutation scheme implemented in fgsea and Benjamini–Hochberg *p*-value adjustment. Genes were ranked by the DESeq2 Wald statistic with no pre-filtering by nominal or adjusted *p*-value, and enrichment direction and magnitude were summarised by normalized enrichment scores (NES) and visualized using the enrichplot package. Genes exhibiting constitutive differential expression attributable to background genomic variation in the *nilr1* accession, unrelated to *NILR1* function, were excluded from the ranking prior to enrichment. For the Gene Ontology (GO) Biological Process analysis of the constitutive genotype effect (Fig. 4f), genes were ranked by the Wald statistic of the pooled baseline contrast (*nilr1* versus wild type under mock; see *Generation of the pooled baseline dataset*) and enrichment was computed with the gseGO function (org.At.tair.db, keyType = “ENTREZID”, ont = “BP”), with minimum and maximum gene-set sizes of 10 and 500 and Benjamini–Hochberg adjustment at adjusted *P <* 0.05.

### Comparisons with published ascaroside, brassinosteroid, and nematode reference sets

The ascaroside-responsive gene set from Letia et al., 2025 was analyzed by the same overlap approach, and rank-based GSEA on the genotype × treatment interaction Wald ranking used clusterProfiler::GSEA (gene set sizes 10–500). A 46-gene reference set spanning BR biosynthesis, core signaling components, and canonical BR-responsive regulators was assembled from Nolan et al., 2020 and intersected with the genotype × treatment interaction tables (Supplementary Tables S5, S6) at each timepoint. The full curated set, with locus identifiers, functional category, and per-gene interaction statistics, is provided in Supplementary Table S7.

For Fig. 5d, post-invasion expression values from Siddique et al., 2022 were matched to NILR1-dependent candidates by Entrez ID, and per-stage log_2_ fold changes computed as log_2_((mean infected + 1) / (mean control + 1)). Pre-invasion values (30 min, 3 h, 10 h) were taken from the DESeq2 outputs of this study.

### Cell-type-resolved expression analysis

Cell-type-resolved expression of *NILR1* (AT1G74360) and *FLS2* (AT5G46330) was extracted from the wild-type Arabidopsis root single-cell atlas of Shahan et al., 2022, accessed via the ARVEX portal (https://shiny.mdc-berlin.de/ARVEX/). Per-cell normalized expression values and cell-type / developmental-zone annotations were used as provided. For each gene, expression was summarized per (cell type × developmental zone) group as the mean normalized expression (AvgExp) and the fraction of cells with detectable expression (PctExp, expression *>* 0).

### Integration with cortex regulatory networks and GTL1/DF1 target spaces

To identify transcription factors (TFs) within the NILR1-dependent candidate sets, the *A. thaliana* TF list was downloaded from the Plant Transcription Factor Database (PlantTFDB v5.0; Jin et al., 2017; https://planttfdb.gao-lab.org/download.php). TAIR locus identifiers from PlantTFDB were mapped to Entrez Gene IDs using org.At.tair.db, and NILR1-dependent candidates at each timepoint were intersected with the resulting TF reference set.

NILR1-associated TFs were mapped to brassinosteroid-responsive cortex elongation GRNs from Nolan et al., 2023 Supplementary Data S4 using Entrez ID matching (via org.At.tair.db). *GTL1* targets were extracted from Supplementary Data S5. Of the 89 (30 min) and 290 (3 h) NILR1-dependent candidates, 77 and 282 respectively had TAIR identifiers in org.At.tair.db and were used for overlap analyses. Two sets were defined: (1) all predicted targets; (2) high-confidence top 30% by absolute regression coefficient. GSEA was performed as described above using clusterProfiler::GSEA, with minimum and maximum gene set sizes of 10 and 5,000. Leading-edge genes and full enrichment results are provided in Supplementary Tables S10 and S11.

Regulatory space membership was defined from published transcriptomic datasets of *df1*, *gtl1*, and *gtl1 df1* mutants (Shahan et al., 2022; Nolan et al., 2023) using an adjusted *P <* 0.05 threshold. Within each regulatory space, infection-induced WT log_2_ fold changes (infected vs. control) were compared between NILR1-dependent candidates and the remaining genes using two-sided Mann– Whitney *U* tests. The resulting *P* -values from all six comparisons (three regulatory spaces × two timepoints) were adjusted for multiple testing using the Benjamini–Hochberg procedure.

### qRT-PCR analysis

Total RNA was reverse-transcribed in 20 µl reactions using the RevertAid First Strand cDNA Synthesis Kit (Thermo Fisher Scientific) according to the manufacturer’s instructions. Quantitative realtime PCR (qRT-PCR) was performed using Fast SYBR Green Master Mix (Applied Biosystems) on a StepOnePlus Real-Time PCR System (Applied Biosystems). Gene-specific primers are listed in Supplementary Table S8. Ct values were normalized to TIP41-like family protein (AT4G34270) to obtain ΔCt values. Statistical analyses were performed on ΔCt values using Welch’s two-sided *t*-tests (within-genotype comparisons) and two-factor linear models (genotype × treatment interactions). For visualization, normalized expression was displayed as 2*^−^*^ΔCt^.

### Apoplastic ROS measurements

Apoplastic reactive oxygen species (ROS) were measured by a luminol-based assay in leaf discs of *A. thaliana* Col-0 and *nilr1*. Discs (3 mm diameter) from 12-day-old seedlings were floated in white 96-well plates with 200 µl water and incubated overnight in the dark to dissipate wound-induced ROS. The water was then replaced with a 100 µl reaction mixture of 35 µl L-012 luminol derivative (8-amino-5-chloro-7-phenylpyrido[3,4-*d* ]pyridazine-1,4-(2*H*,3*H*)dione sodium salt; FUJIFILM Wako, Japan), 15 µl horseradish peroxidase (HRP; 20 µg ml*^−^*^1^) and 50 µl treatment solution. Luminescence was recorded immediately at 80 s intervals for 90 min on a TECAN Infinite 200 Pro reader (Tecan) as relative light units (RLU). Each treatment had at least twelve biological replicates. Cumulative ROS (area under the curve, AUC) was analyzed per replicate by two-way ANOVA with genotype and treatment as fixed factors, followed by Tukey’s HSD and compact letter displays at *α* = 0.05.

## Results

### NILR1 contributes to quantitative resistance against *H. schachtii*

Mendy et al., 2017 showed that *nilr1* loss-of-function mutants display increased susceptibility to *H. schachtii*. To characterise this phenotype under our experimental conditions, we quantified parasitism success in Col-0 and *nilr1* mutants. At 12 days post infection (dpi), *nilr1* mutants showed a 23.7% increase in male numbers, and a 16.6% increase in total nematode numbers compared with wild type (*P* = 0.004, *P* = 0.028, respectively), whereas female numbers were unchanged (Fig. 1a, *P* = 0.42). At 14 dpi, female nematodes were significantly larger by 5.8% in *nilr1* than in wild type (Fig. 1b, *P* = 0.010), whereas syncytium size did not differ significantly between genotypes (Fig. 1c, *P* = 0.074). These observations confirm a reproducible, statistically supported shift towards higher nematode burden and slightly increased female size in *nilr1* under our assay conditions, consistent with a contributory role of NILR1 in quantitative resistance to *H. schachtii*.

### NILR1 is a cortex-expressed BRI1-clade receptor induced by the general stress regulon

To investigate how the NILR1 receptor kinase influences early transcriptional responses to *H. schachtii* infection, we performed a comparative RNA-seq analysis of 12-day-old wild-type (Col-0) and *nilr1* mutant *A. thaliana* roots following inoculation with *H. schachtii* infective second-stage juveniles (J2s). Root tissues were harvested at 30 min and 3 h post inoculation, alongside uninfected controls (Fig. 2a), to capture early and intermediate phases of the host response.

In wild-type roots, nematode infection triggered a moderate transcriptional response at 30 min post inoculation, with 417 upregulated and 39 downregulated genes (adjusted *P* ≤ 0.05, | log_2_ FC| ≥ 1), expanding to 2726 upregulated and 867 downregulated genes by 3 h (Fig. 2b, Tab. S1-S4). *nilr1* mutants showed a comparable overall response at both time points (350 up / 92 down at 30 min; 2643 up / 1026 down at 3 h), but with a consistent shift toward transcript downregulation and reduced induction relative to wild type. At baseline, 1340 genes were de-repressed and 852 were repressed in *nilr1* mutants, compared to WT (Fig. 2c, Tab. S12). NILR1 was the only BRI1-clade (LRR-RLK-Xb) receptor kinase strongly induced at 30 min, with the largest induction amplitude across the clade at 3 h (Fig. 2d), showing a pronounced ≈16-fold induction (log_2_ FC ≈ 4) during the early perception phase, with transcript levels remaining elevated at 3 h and returning to near-baseline at post-invasion stages (Siddique et al., 2022, Fig. 2e).

Re-analysis of published datasets (Fig. 2f) indicated that NILR1 transcript abundance is also increased in Arabidopsis roots and shoots in response to diverse biotic and abiotic stimuli—including bacterial and oomycete infection, elicitor peptides, chitin, oligogalacturonides, wounding and ionic/osmotic stress and mildly antagonised by ABA signaling, placing NILR1 among a broadly stress-responsive transcriptional cohort in these experiments (Mendy et al., 2017; Safaeizadeh et al., 2024; Ramirez et al., 2026; Stringlis et al., 2018; Davidsson et al., 2017; Rymen et al., 2019; Bjornson et al., 2021; Lamers et al., 2025). Single-cell transcriptomic data from wild-type roots (Shahan et al., 2022) showed the highest abundance of *NILR1* in cortex cells, with additional expression in endodermis, epidermis, and procambium (Fig. 2g), spanning the layers engaged during nematode approach and cortex migration. In the same atlas, FLS2 transcripts were detectable but displayed lower mean expression and a more restricted pattern, with prominent signals in atrichoblasts of the maturation zone and in xylem-adjacent cells, suggesting partly non-overlapping spatial distributions of these receptors (Fig. S4).

### Canonical PTI marker responses are preserved in *nilr1* during early nematode infection

To determine whether NILR1 is required for activation of canonical PTI during early nematode infection, we evaluated expression levels from the RNAseq data. Canonical PTI marker genes, including *WRKY29*, *WRKY33*, *FRK1*, *GSTF6*, and *PR4*, were robustly induced by nematode infection in both genotypes in our RNA-seq data and qRT–PCR assays (Fig. 3a,b). Where the genotype × treatment interaction reached significance, the effect consistently reflected *elevated* rather than diminished transcript accumulation in infected *nilr1* roots: this was the case for *FRK1* and *CYP81F2* by qRT–PCR (Fig. 3b), indicating that the markers tested here are not attenuated by loss of NILR1.

**Figure 3:**
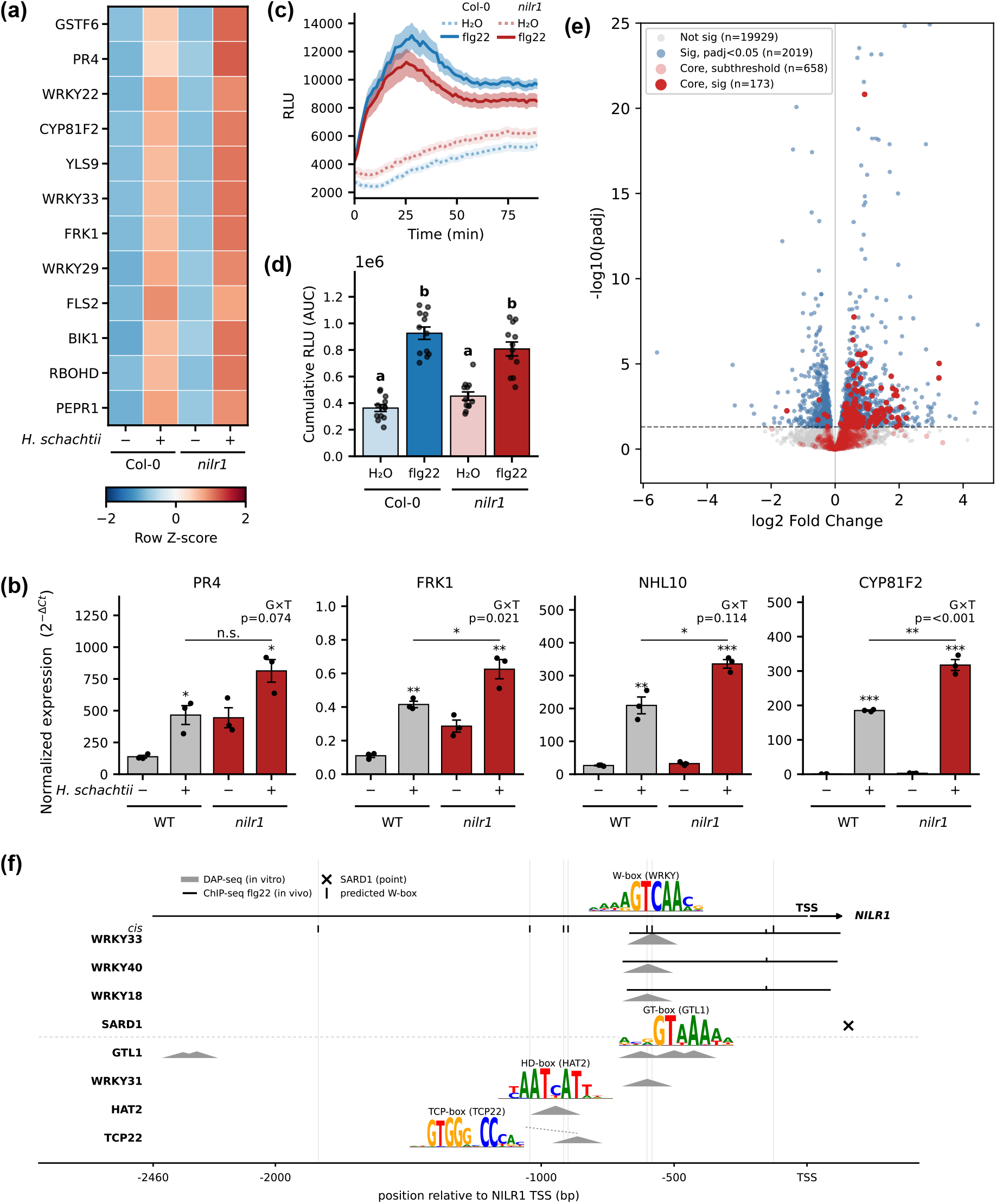
Canonical PTI-associated transcription is induced during early nematode infection independently of *NILR1*. **(a)** Heatmap of canonical PTI marker gene expression in Col-0 and *nilr1* roots at 3 h post inoculation with *H. schachtii*. Row Z-scores of normalized RNA-seq counts (*n* = 4). **(b)** Expression of canonical PTI marker genes in Col-0 and *nilr1* roots at 3 h post inoculation with *H. schachtii*, assessed by qRT–PCR. Bars represent the mean of three biological replicates (*n* = 3) ± SE. Asterisks above bars indicate within-genotype significance and brackets compare WT vs *nilr1* under infection (Welch’s two-sided *t* -test on Δ*Ct*): *^∗^P <* 0.05, *^∗∗^P <* 0.01, *^∗∗∗^P <* 0.001. G×T *P*-values indicate the genotype × treatment interaction. **(c)** ROS production in Col-0 and *nilr1* leaf discs treated with 0.5 µM flg22 or water, recorded over 90 min. Traces show mean relative light units (RLU); shaded areas indicate ± SEM (*n* = 12 leaf discs per treatment). **(d)** Cumulative ROS production over 90 min from the traces in (**c**). Bars show mean ± SEM; dots represent individual leaf discs (*n* = 12). Letters denote Tukey HSD groups following two-way ANOVA with genotype and treatment as fixed factors (*p <* 0.05). Representative of three independent experiments. These data indicate that the PTI markers and flg22-triggered ROS outputs assayed here are not reduced in *nilr1*. **(e)** Volcano plot of baseline transcriptional differences (*nilr1* vs wild type, log2FC *>* 0, adjusted *P <* 0.05, *n* = 8), with core PTI genes (Bjornson et al., 2021) highlighted; the set is strongly enriched among genes de-repressed at rest in *nilr1*, showing that NILR1 constitutively restrains the PTI program in the uninfected root. **(f)** Transcription-factor occupancy across the *NILR1* promoter, relative to the TSS. Grey triangles, *in vitro* DAP-seq peaks (O’Malley et al., 2016); black bars, *in vivo* flg22 ChIP-seq–bound regions (Birkenbihl et al., 2017); cross, SARD1 site (T. Sun et al., 2015); logos, consensus W-box, GT-box, HD-box and TCP-box.

In luminol-based ROS assays with leaf discs, flg22-triggered ROS kinetics, peak amplitudes and cumulative output (AUC) did not differ significantly between Col-0 and *nilr1* (AUC; Tukey-adjusted *P* = 0.18; *n* = 12 per treatment; Fig. 3c,d, S5), suggesting that this canonical FLS2/BAK1/RBOHD-dependent output is intact in the mutant under these conditions.

Among 1340 genes with significantly higher basal expression in *nilr1* than in wild type (baseline log_2_FC *>* 0, adjusted *P <* 0.05 vs. WT under mock), core PTI genes (Bjornson et al., 2021) were strongly over-represented (167 observed vs 48.9 expected by chance; 3.4-fold enrichment; one-sided hypergeometric test, *P* = 1.6 × 10*^−^*^46^), whereas only 6 core PTI genes were significantly downregulated at baseline. This de-repressed set included classical PTI and defense markers (e.g. *FRK1*, *MPK3*, *CYP71A12*, *GSTF7*, *WAKL10*) as well as components of the Pep danger signaling module, including the receptor PEPR2 and the Pep-peptide precursor PROPEP3 (alongside PROPEP1/4/5/6).

#### The *NILR1* promoter is under immune and developmental transcriptional control

Because *NILR1* was strongly infection-induced yet dispensable for canonical PTI output, we examined how *NILR1* itself is transcriptionally regulated (Fig. 3f). Its promoter carries a cluster of W-boxes ∼580–600 bp upstream of the TSS and an additional W-box near −126. *In vivo*, *NILR1* was a direct target of the core immune regulators *WRKY18*, *WRKY40* and *WRKY33* following flg22 elicitation (Birkenbihl et al., 2017) and of the immune master regulator *SARD1* (T. Sun et al., 2015). *In vitro* DAP-seq independently placed *WRKY33/40/18* peaks over the same −583 cluster (O’Malley et al., 2016). Consistent with this, *WRKY18*, *WRKY40*, *WRKY33* and *SARD1* were themselves rapidly and strongly induced by nematode infection in both wild-type and *nilr1* roots.

Four of the promoter-binding factors identified by DAP-seq — *GTL1*, *WRKY31*, *HAT2* and *TCP22* —were also among the NILR1-dependent candidate TFs recovered from the interaction analysis; we therefore asked whether each occupies a genuine cognate element. Re-scanning every factor’s own DAP-seq matrix against its peak confirmed the expected site for three of the four: the GT-element beneath *GTL1* (−500), a homeodomain TAAT/ATTA motif beneath *HAT2* (−946), and a canonical Wbox beneath WRKY31 (−601). The *WRKY31* peak, however, coincides with the same −583/−601 W-box cluster bound by the immune WRKYs and thus adds no independent cis-element; *WRKY31* was itself infection- and baseline-induced, consistent with feedback within the WRKY regulon. The *TCP22* peak (−864) carried only a weak, sub-consensus TCP element. Thus, the *NILR1* promoter combines at least two regulatory codes: an immune module that drives its infection induction, and a developmental module whose factors putatively set its expression.

We therefore interpret NILR1 primarily as a negative regulator of basal PTI gene expression in the uninfected root, rather than as an essential activator of the canonical PTI markers analyzed here. From a purely transcriptional perspective, the elevated basal PTI signature in *nilr1* might be expected to favour resistance; however, the observed increase in *H. schachtii* parasitism indicates that this constitutive immune shift alone is not sufficient to confer enhanced protection in our system. Accordingly, while the basal immune phenotype likely reflects one aspect of NILR1 function, it appears unlikely to be the sole determinant of the quantitative resistance phenotype we observe under nematode infection.

### NILR1-sensitive differences in the early infection response

To resolve genotype-specific differences in the early infection response, we applied a genotype × treatment interaction model (∼genotype + treatment + genotype : treatment) comparing wild-type and *nilr1* roots, which tests whether the transcriptional response to *H. schachtii* differs between genotypes independently of basal expression. In this framework, the interaction term captures genes whose infection response differs between wild type and *nilr1* after accounting for main effects. Because interaction terms are estimated with lower power than main effects, we used the apeglm shrinkage estimator to obtain stabilised effect-size estimates (shrunken log2 fold-changes) and prioritised genes with interaction estimates ≤ -0.2 as NILR1-sensitive candidates for downstream analysis, irrespective of formal FDR significance. Throughout, we therefore interpret these genes as showing NILR1-sensitive infection responses in our model, rather than as strictly NILR1-dependent in a mechanistic sense.

At 30 min post infection, 4 of 25,278 tested genes reached formal significance (padj ≤ 0.05) for the interaction term, reflecting the expected low power of interaction tests at this early time point. Shrinkage-based prioritisation identified 89 NILR1-dependent candidates and 44 enhanced responses (Fig. 4a). By 3 h post infection (Fig. 4b), interaction effects became more pronounced at the whole-transcriptome level—10 of 25,453 tested genes reached formal significance, nine with negative interaction statistics—yielding 290 NILR1-dependent candidates and 179 enhanced responses, a progressive bias toward reduced infection-induced activation in *nilr1*. We therefore use the shrinkage-prioritised sets as a structured ranking for enrichment and network analyses, while interpreting individual-gene effects cautiously.

**Figure 4:**
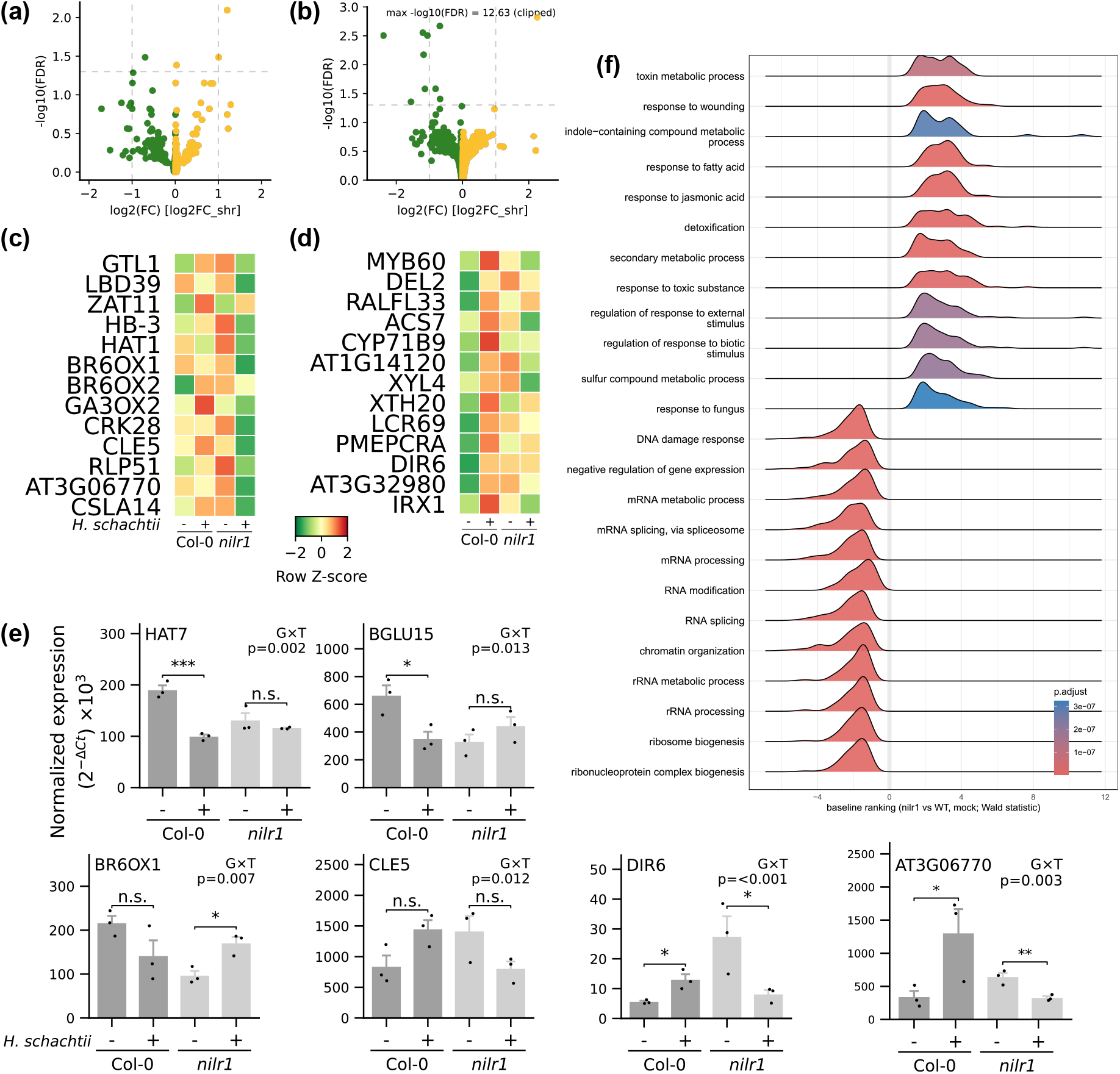
**(a,b)** Volcano plots of genotype × treatment interaction effects at 30 min (**a**) and 3 h (**b**). Interaction log2 fold changes represent shrunken effect-size estimates of NILR1-dependent differences in nematode-induced gene expression between Col-0 and *nilr1*, with negative values indicating attenuated responses in the mutant. **(c,d)** Row-scaled expression (Z-scores of normalized counts) of NILR1-dependent candidates at 30 min **(c)** and 3 h **(d)** across wild-type and *nilr1* roots under control and infected conditions. **(e)** Expression of selected NILR1-dependent candidates verified by qRT-PCR in wild-type and *nilr1* roots at 30 min and 3 h post inoculation. Dark grey bars, Col-0; light grey bars, *nilr1*. Within-genotype comparisons (control vs infected) were assessed using Welch’s two-sided *t*-test on Δ*Ct* values. G×T *P* -values indicate the significance of the genotype × treatment interaction (two-factor linear model, *^∗^P <* 0.05, *^∗∗^P <* 0.01, *^∗∗∗^P <* 0.001). Bars represent the mean of three biological replicates (*n* = 3) ± SE. **(f)** Gene set enrichment analysis of the *nilr1* baseline phenotype. Genes were ranked by the Wald statistic of the baseline contrast (*nilr1* vs wild type, mock), with positive values indicating de-repression in *nilr1*, and rank-based GSEA was performed for Gene Ontology Biological Process categories. Each ridge shows the distribution of ranks of a term’s leading-edge genes; fill color indicates the adjusted *P* value and terms are ordered by normalized enrichment score. Only terms with adjusted *P <* 0.05 are shown. Gene sets attenuated in *nilr1* encompass mRNA splicing and processing, RNA modification, chromatin remodeling, ribosome biogenesis, and ribonucleoprotein complex assembly. Gene sets enhanced in *nilr1* are dominated by glucosinolate and secondary metabolism, jasmonic acid and wounding responses, detoxification, and antifungal defense categories.

By construction, genes whose expression differs between genotypes to a similar extent under mock and infected conditions have interaction coefficients close to zero. In our data, 101 of 379 shrinkage-prioritised candidates (27%) also showed significant baseline differences (adjusted *P <* 0.05 under mock), indicating that the majority of NILR1-sensitive infection effects arise on top of, rather than being trivially explained by, constitutive genotype differences.

At 30 min, the NILR1-sensitive gene set was enriched for transcription factors associated with developmental regulation, including HD-ZIP factors (HAT1, HAT2, HB20), GTL1 and cell-cycle/patterning regulators (E2F3, MYB66), whereas classical immune families such as WRKY and NAC were not recovered at this early time point (Fig. 4c,e). By 3 h, the TF composition shifted towards bHLH, MYB, C2H2 and TCP families, which include factors previously implicated in stress and defense, while developmental regulators remained represented, suggesting that developmental and defense-associated regulatory components become progressively intertwined during early infection (Fig. 4d,e, Table S16). The strongest transcription-factor effect was the R2R3-MYB factor *MYB60* (shrunken log_2_FC = −1.21; qRT-PCR validated, Fig. S8). Notably, the bHLH factor *NAI1* and its known target *PYK10* (Matsushima et al., 2004) were co-regulated within the same set, placing regulator and output inside a single NILR1-sensitive module, and *WRKY31* was the only canonical defense-associated TF present.

The NILR1-sensitive candidate set at this timepoint also encompassed several functionally coherent groups of genes, spanning successive steps of cell-wall biology (expansins, hemicellulose/matrix remodeling, lignin-associated components), indole glucosinolate metabolism from biosynthesis to hydrolysis, and peptide/hormone signaling (e.g. PEPR2, ADR1/ADR1-L2, RALF1/RALFL33, ethylene/auxin/jasmonate enzymes, Fig. 4c,d). The breadth of pathway coverage within these groups suggests that NILR1-sensitive infection responses extend across multiple layers of cell wall, secondary metabolism and signaling, rather than being confined to isolated single genes.

We therefore treat the candidates as NILR1-dependent differences in the infection response and characterise that resting state directly in the following section.

### Loss of NILR1 is associated with a growth-to-defense shift at baseline

To characterise the basal transcriptome differences between genotypes, we performed GSEA on *nilr1* versus Col-0 under mock conditions (Fig. 4f, Supplementary Table S12). Gene sets elevated in *nilr1* were dominated by defense- and secondary-metabolism categories (jasmonate signaling and response, glucosinolate and indole/camalexin biosynthesis, glutathione-dependent detoxification), whereas sets reduced in *nilr1* were enriched for growth-associated biosynthetic machinery (ribosome biogenesis, rRNA processing, RNA splicing, chromatin organisation). Thus, at the whole-root level and under our growth conditions, loss of NILR1 is associated with a shift in baseline transcriptional investment away from growth-related gene sets and towards defense- and stress-associated gene sets.

Within this baseline shift, we could distinguish modules with elevated expression in *nilr1* (e.g. a jasmonate-associated module and a lignin/secondary-wall module) and a cytokinin-associated module with reduced expression. None of these modules showed significant enrichment among genotype × treatment interaction effects, indicating that they respond to nematode infection in broadly similar ways in both genotypes. Accordingly, the growth-to-defense bias appears to be established prior to infection and then largely propagated through the early infection time course, rather than being generated or resolved by the infection itself in a NILR1-specific manner.

Taken together, these observations indicate that, at baseline and for the defense-related gene sets examined here, loss of NILR1 is not associated with a reduction but rather with an overall increase in the transcriptional representation of multiple defense and secondary-metabolism pathways. Under infection, these pathways remain inducible in both genotypes. The elevated defense signature therefore makes it unlikely that the increased susceptibility of *nilr1* is caused by a simple deficit in the broad defense programs captured by our transcriptome-level GSEA, although we cannot exclude more specific or post-transcriptional immune alterations. Instead, the transcriptional shift towards defense coincides with a growth-reduced state, consistent with previous reports of shorter primary roots and reduced cell expansion in *nilr1* (*grace-1*) plants (Z. Wu et al., 2017; Zheng et al., 2022); this suggests that altered growth–defense trade-offs, rather than immune failure per se, may contribute to the quantitative susceptibility phenotype, a possibility that will require direct experimental testing.

### NILR1 signals via a brassinosteroid-type kinase cascade, while nematode-induced BR transcriptional responses remain largely NILR1-independent

Previous work has placed NILR1 within a brassinosteroid-type receptor module that includes the coreceptor BAK1 and BSK substrates (Mendy et al., 2017; Zheng et al., 2022). In line with this, our data confirm transcriptional induction of NILR1 during infection but do not provide evidence for a major NILR1-dependent reconfiguration of the canonical BR transcriptional program under nematode attack. Among a curated set of 46 BR biosynthesis genes, core signaling components, and BES1/BZR1 targets, only the late biosynthetic enzymes *BR6OX1* (*CYP85A1*) and *BR6OX2* (*CYP85A2*) showed NILR1-dependent interaction effects at 30 min, and no genes did at 3 h (Supplementary Fig. S9, Table S7). Infection coordinately downregulated multiple BR biosynthetic enzymes (BR6OX1, DWF1, DWF4) and BES1 while inducing the negative regulator BIN2 — a response consistent with infection-associated suppression of BR signaling. These parallel infection responses in wild type and *nilr1* suggest that, at least at the early time points and for the BR-related gene sets examined here, the broader BR transcriptional response to *H. schachtii* is largely NILR1-independent.

### NILR1-sensitive transcription factors map onto a HAT7/GTL1 cortex developmental network

To determine whether NILR1-dependent transcription factors converge on an established developmental regulatory architecture, we mapped the TFs identified at 30 min (14) and 3 h (16) onto the brassinosteroid-responsive cortex elongation GRN of Nolan et al., 2023, in which *HOMEOBOX FROM ARABIDOPSIS THALIANA 7*(*HAT7*) ranked as the top hub regulator and *GT-2-LIKE 1* (*GTL1*) among the top five TFs by network centrality. Thirteen of 14 TFs at 30 min (93%) and 11 of 16 at 3 h (69%) were present in this network (Fig. 5a). The single TF outside the network at 30 min was *LBD39*; at 3 h, five TFs mapped outside, including the R2R3-MYB factor *MYB60*, which binds the promoters of several secondary cell wall genes, including *CESA8*/*IRX1*, in yeast one-hybrid assays (Taylor-Teeples et al., 2015). Thus, most NILR1-sensitive TFs fall within a previously defined cortex developmental network, consistent with NILR1 affecting infection-induced transcription through TFs that are already wired into this BR-responsive developmental module.

**Figure 5:**
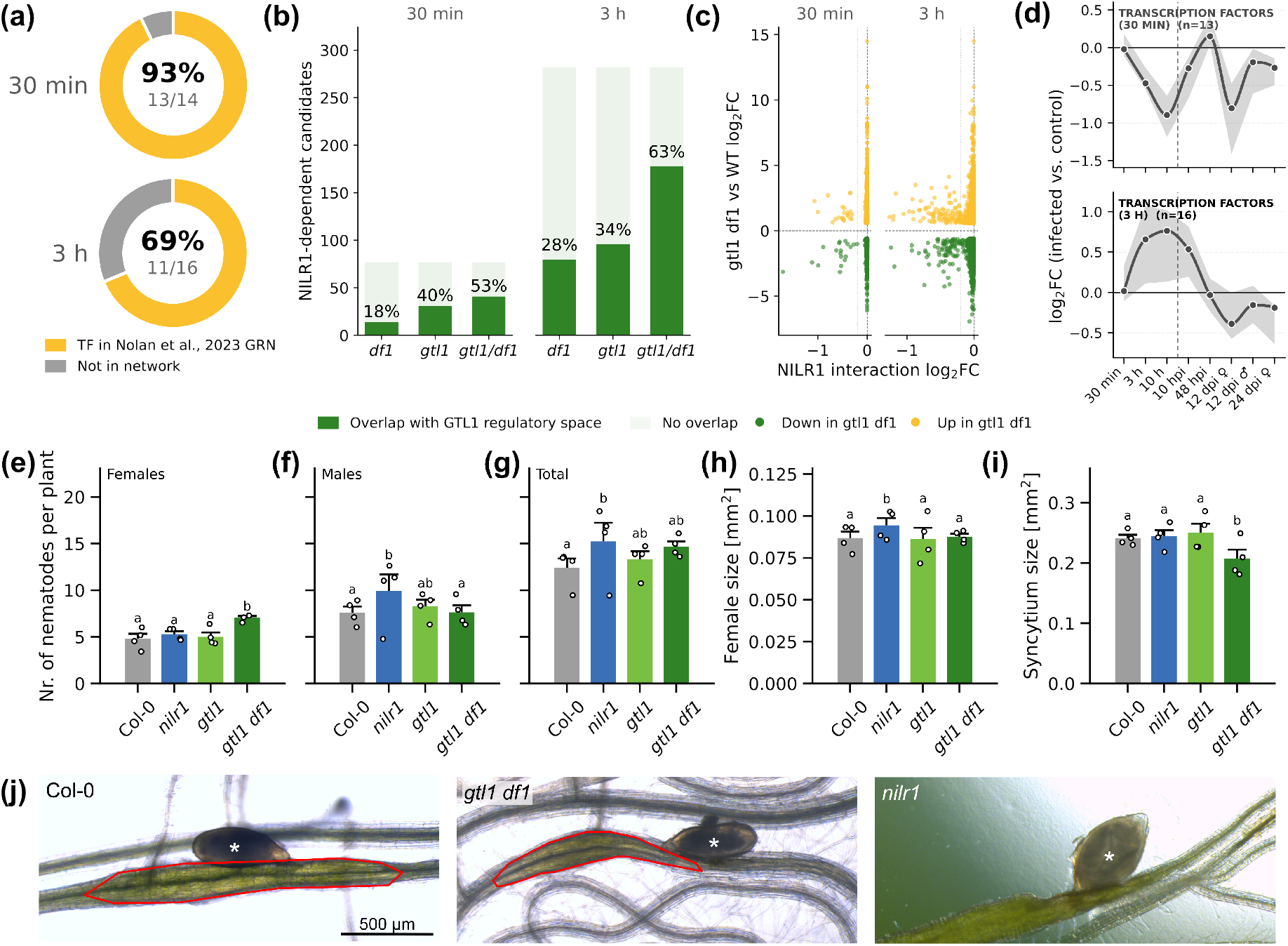
NILR1-dependent transcription factors map to a cortex-enriched regulatory network. **(a)** Proportion of NILR1-dependent transcription factors present in the brassinosteroid-responsive cortex elongation GRN (Nolan et al., 2023) at 30 min and 3 h post inoculation. **(b)** Overlap between NILR1-dependent candidate genes (interaction log2FC *<* −0.2) and transcription factor regulatory spaces defined by genes misregulated in *df1*, *gtl1*, and *gtl1 df1* mutants. **(c)** Relationship between NILR1-dependent transcriptional responses and the *GTL1*/*DF1* regulatory space at 30 min and 3 h following *H. schachtii* inoculation. Scatter plots show *NILR1* genotype × treatment interaction effects plotted against transcriptional responses observed in the *gtl1 df1* mutant relative to WT. Colors indicate genes up-or downregulated in *gtl1 df1*. **(d)** Temporal trajectories of NILR1-dependent gene modules in Col-0 roots across early perception (30 min, 3 h, 10 h; this study) and post-invasion stages (10 hpi–24 dpi; Siddique et al., 2022), shown as log_2_ fold change (infected vs. control). Bold lines indicate the per-timepoint median across genes within each module, shaded ribbons the 25th–75th percentile range, and faint traces individual genes. Analysis of the *H. schachtii* infection phenotype on the *gtl1* single mutant and the *gtl1 df1* double mutant, compared to *nilr1* and WT. The number of **(e)** female, **(f)** male and **(g)** total *H. schachtii* per plant was evaluated at 12 dpi. Female **(h)** and associated syncytium **(i)** sizes were measured at 14 dpi. Barplots show the average of *n* = 4 independent biological replicates; circles represent individual replicates. Letters above groups denote significant differences at *α* = 0.05 (S^̌^ idák-adjusted compact letter display); groups sharing a letter are not significantly different. See Methods for full statistical models and pairwise tests. **(j)** Representative stereomicroscope images of *H. schachtii* females and associated syncytia at 14 dpi in Col-0, *gtl1 df1*, and *nilr1* roots. The female nematode is indicated by the white asterisk; the red outline indicates the nematode-induced feeding site (syncytium). Images illustrate the genotype-specific phenotypes quantified in (h,i): enlarged females in *nilr1* and reduced syncytium size in *gtl1 df1*, both relative to Col-0.

*GTL1* showed the strongest interaction effect of any TF at 30 min, and *HAT7*, the top-ranked hub of the Nolan et al. GRN, was independently confirmed by qRT-PCR (Fig. 4e, G×T p=0.002). The 30 min set additionally included a cluster of HD-ZIP regulators (*HAT1*, *HAT2*, *HB-2*, *HB20*) that are neighbours of *HAT7* and *GTL1* within the same network.

Because *GTL1* ranked highest among NILR1-dependent TFs at 30 min, we asked whether genes regulated by *GTL1* and its homolog *DF1* (Shibata et al., 2018; Nolan et al., 2023, Supplementary Table S9) were enriched among NILR1-dependent responses. GSEA of the genotype × treatment interaction ranking revealed significant enrichment of predicted *GTL1* targets toward negative interaction statistics, strengthening from 30 min to 3 h for both the high-confidence target set (NES = −1.85 → −1.99; adjusted *P* = 3.5 × 10*^−^*^4^ → 9.5 × 10*^−^*^6^; Supplementary Fig. S10) and the inclusive set of 294 targets (NES = −1.76 → −1.89; adjusted *P* = 1.7 × 10*^−^*^6^ → 2.5 × 10*^−^*^8^; Supplementary Table S10). Leading-edge genes are listed in Supplementary Table S11. This enrichment indicates that genes previously placed in the GTL1/DF1 regulatory space are disproportionately represented among genes whose nematode-induced responses are attenuated in *nilr1* in our dataset.

Comparing strongly NILR1-sensitive candidates with genes misregulated in *gtl1 df1* mutants (Nolan et al., 2023) showed that 41 of 77 candidates (53%) at 30 min and 178 of 282 (63%) at 3 h fall within the GTL1/DF1 regulatory space, whereas *gtl1 df1* misregulates 8,391 genes overall (Fig. 5b). This suggests that NILR1-sensitive infection responses occupy a relatively focused subset of the broader GTL1/DF1-controlled transcriptome. Within this overlap, NILR1-sensitive genes are found among both up- and downregulated fractions of the *gtl1 df1* response (Fig. 5c), indicating that NILR1-associated infection effects intersect both the activating and repressing arms of the GTL1/DF1 module rather than a single polarity. We therefore interpret NILR1 as influencing a specific subspace of the GTL1/DF1 network during infection, rather than globally reprogramming the entire module.

Beyond the GTL1/DF1 module, individual NILR1-dependent candidates included cell-cycle regulators (*E2FE* at 30 min; *DEL2*, *ORC1B*, *CYCD4;1* at 3 h) and a second set of secondary cell wall biosynthetic genes (*GAUT12*, *RWA3*, *IRX1*/*CESA8*; Taylor et al., 2000). Curated cell-cycle target sets (Gombos et al., 2023) showed no coordinated NILR1-dependent behavior, and of the cell wall genes only *IRX1*/*CESA8* is a direct *MYB60* binding target (Taylor-Teeples et al., 2015), with *MYB60* itself repressed at 3 h.

In summary, the HAT7/GTL1 cortex network, probed here via its GTL1/DF1 arm, accounts for a substantial and functionally coherent fraction of the NILR1-sensitive transcriptional changes we observe during early infection, particularly those linked to cell wall and growth-related processes.

Across the combined early (30 min, 3 h, 10 h) and post-invasion infection stages (10 h–24 dpi) in wild type, the two TF sets showed opposing temporal profiles: the 30 min set (*n* = 13; HAT7, GTL1 and HD-ZIP neighbours) was transiently suppressed, reaching log_2_FC ≈ −1 at 10 h before recovering by 48 hpi, whereas the 3 h set (*n* = 16; bHLH, MYB, C2H2, TCP) was induced at 3 h (log_2_FC ≈ +0.8) and remained elevated at 10 h. These opposing trajectories are consistent with an early, transient downregulation of growth-associated hubs followed by sustained induction of TFs previously linked to stress and defense, in line with a general growth–defense adjustment during the infection time course.

#### Genetic disruption of the GTL1/DF1 module alters *H. schachtii* parasitism

The substantial overlap between NILR1-dependent genes and GTL1/DF1 target genes prompted us to evaluate the relevance of the GTL1/DF1 module for *H. schachtii* parasitism. We quantified infection parameters in wild-type, *nilr1*, *gtl1* and *gtl1 df1* plants, scoring females, males and the total number of nematodes per plant (Fig. 5). Consistent with our earlier results, *nilr1* mutants were significantly more susceptible than Col-0, with male and total nematode counts elevated by 37.8% (*P* = 0.002) and 25.6% (*P* = 0.013), respectively, while female numbers were unchanged (*P* = 0.88). The *gtl1* single mutant showed no detectable susceptibility phenotype, with female, male and total counts all indistinguishable from Col-0 (all *P* ≥ 0.77). In contrast, the *gtl1 df1* double mutant displayed a distinct, female-specific profile: female numbers were increased by 46.9% relative to Col-0 (*P* = 0.002), and were likewise elevated above *nilr1* and *gtl1* (*P* = 0.017 and *P* = 0.019), whereas male and total counts remained at wild-type levels (all *P* ≥ 0.26). We next assessed nematode development by measuring female and associated syncytium size (Fig. 5h,i). Female size was increased by 8.1% in *nilr1* relative to Col-0 (*P* = 0.014), while *gtl1* and *gtl1 df1* were indistinguishable from wild type. Conversely, syncytium size was reduced specifically in *gtl1 df1*, by 14.9% relative to Col-0 (*P <* 0.001), with *nilr1* and *gtl1* matching Col-0.

## Discussion

The *Arabidopsis* receptor kinase NILR1 contributes to quantitative resistance to the cyst nematode *H. schachtii*, yet its signaling identity has remained contested: functional and biochemical studies place NILR1 within a conserved brassinosteroid-type kinase cascade associated with growth regulation (Z. Wu et al., 2017; Zheng et al., 2022; Ali et al., 2025), whereas elicitor-based assays have implicated it as a pattern-recognition receptor mediating nematode-associated immune activation (Mendy et al., 2017; Huang et al., 2023, 2024).

Several observations argue against a primary role for NILR1 in canonical PTI: flg22-responsive genes were induced similarly in wild-type and *nilr1* roots, canonical PTI markers remained robustly inducible in the mutant, and flg22-triggered ROS production did not differ significantly between genotypes. Loss of NILR1 is in fact associated with a stronger immune transcriptome at baseline: the core-PTI program (Bjornson et al., 2021) is constitutively de-repressed in uninfected *nilr1* roots (Fig. 3e) yet remains normally inducible under infection. This combination – elevated basal PTI signature together with retained inducibility – makes it unlikely that the increased susceptibility of *nilr1* is caused by a simple deficit in the broad PTI outputs captured by these assays, which would be expected to favour resistance rather than susceptibility.

This functional separation is consistent with NILR1’s signaling architecture: as a BRI1-clade LRR-RLK coupled to a brassinosteroid-type kinase cascade (Z. Wu et al., 2017; Zheng et al., 2022; Ali et al., 2025), NILR1 is not wired to the MAPK output through which LRR-RLK-XII receptors such as FLS2 activate WRKY- and FRK1-type defense transcription (Jones et al., 2024; Ngou et al., 2026). The promoter analysis helps explain how a receptor so tightly connected to defense-related transcription can be dispensable for the canonical PTI outputs we measured: *NILR1* is a direct target of several WRKY immune regulators and SARD1 (Birkenbihl et al., 2017; T. Sun et al., 2015; Fig. 3f), so its infection induction—and its induction across every biotic and abiotic stress we examined (Fig. 2f)— is an output of the immune transcriptional network, not an input to it. NILR1 sits downstream of PTI, switched on by the same WRKY/SARD1 machinery it is dispensable for, while its own output converges on the HAT7/GTL1 developmental module. It is thus embedded in immunity yet decoupled from it: PTI-related transcription drives *NILR1*, but NILR1 does not drive PTI.

This bears directly on how NILR1’s ligand is understood. The prevailing interpretation, based largely on exogenous applications, casts Ascr18 as a nematode-derived molecular pattern perceived by NILR1 to activate immunity (Huang et al., 2023), including NILR1-dependent MPK3/MPK6 activation and resistance in certain pathosystems. In such a model, loss of NILR1 would be expected to weaken canonical immunity. In contrast, our data show that the canonical PTI markers and flg22-triggered ROS outputs we examined are preserved or even constitutively elevated in *nilr1*, arguing against a scenario in which NILR1 is generally required for these particular arms of PTI in roots. Three additional observations highlight that the relationship between Ascr18 perception and NILR1’s function in our system is likely to be more nuanced than a simple *immune sensor* model. First, the published Ascr18-responsive transcriptional programme (Letia et al., 2025) overlaps only weakly with the NILR1-dependent component of the nematode infection response in our early root assays (Fig. S6, S7). Second, purified Ascr18 alone did not strongly induce classical *PR1*, *FRK1*, or *WRKY* markers in the recent priming study of Manohar et al., 2026, but instead potentiated their pathogen-triggered expression, indicating a prominent role in priming rather than in direct activation. Third, it remains unknown whether *H. schachtii* delivers Ascr18 at physiologically relevant concentrations *in planta* (Yang et al., 2026), so current *in vitro* binding data (Huang et al., 2023) establish affinity but do not yet define the *in situ* contribution of Ascr18 to NILR1 signaling during infection. We therefore propose that the quantitative resistance phenotype and transcriptional patterns described here can be explained without invoking a primary, indispensable role of Ascr18-triggered PTI in this particular root–nematode interaction. Because the PRR–MAPK immune arm appears fully preserved, pathogen sensing via canonical PRRs is likely NILR1-independent in our system. An attractive – but at present speculative – possibility is that, during early root infection, NILR1 predominantly reads one or more endogenous cues, for example stress- or developmentally regulated signals, and that nematode-derived ascarosides modulate this endogenous signaling state rather than acting as the sole trigger. The physiologically relevant spectrum of NILR1 ligands thus remains undefined and will require direct ligand-delivery and genetic tests.

Nolan et al. (2023) identified HAT7 and GTL1 as top-ranked brassinosteroid-responsive transcription factors in the elongating cortex gene regulatory network, where they promote cell elongation and cell wall–related gene expression. Both are direct BES1/BZR1 targets (Y. Sun et al., 2010; Yu et al., 2011; Oh et al., 2014), and GTL1 physically interacts with BES1 to co-regulate a shared set of brassinosteroid-induced targets in the cortex (Nolan et al., 2023). Our NILR1-sensitive transcriptome engages multiple components of this architecture: at 30 min, the NILR1-dependent TF set includes *GTL1* and *HAT7* themselves, along with additional HD-ZIP regulators. *GTL1* and *DF1* have been implicated across diverse developmental and stress contexts (Breuer et al., 2009; Shibata et al., 2018, 2022; Vö lz et al., 2018; Mano et al., 2024; Chen et al., 2025; Urzúa Lehuedé et al., 2025), positioning them as context-dependent regulatory hubs whose output is shaped by upstream input and tissue context. To our knowledge, NILR1 has not previously been placed within this regulatory architecture: Zheng et al., 2022 showed that NILR1 activates the BR-type kinase cascade, sharing the co-receptor BAK1 and BSK substrates with BRI1 and driving BES1 dephosphorylation, while Nolan et al., 2023 identified HAT7 and GTL1 as direct BES1/BZR1 targets. Our data add an infection-context layer by showing that loss of NILR1 selectively alters the transcriptional output of a defined subset of this module under nematode challenge.

Notably, the two NILR1-sensitive TF sets with opposite temporal behaviour—the early-suppressed HAT7/GTL1 growth hubs and the later-induced defense regulators—map predominantly to the same cortex network. This indicates that NILR1 shapes the output of a single developmental module as it transitions from growth to defense during early infection, rather than acting on scattered targets.

Whether NILR1-associated transcriptional changes during early infection are mounted in cortex cells themselves or in surface-layer cells that redeploy a cortex-characterised GRN cannot be resolved from bulk root transcriptomes; cell-type-resolved profiling will be required to localise the resistance-relevant response. Likewise, the downstream consequences that this signature predicts – altered cortex cell-wall properties and glucosinolate composition – remain to be tested by targeted biochemical and biomechanical assays.

A regulatory cascade runs from receptor to downstream target: the ER-body *β*-glucosidase *PYK10* is not a direct GTL1 target but is activated by the bHLH factor NAI1 (Matsushima et al., 2004), which is itself both NILR1-dependent and embedded within the GTL1/DF1 network — placing the full regulatory chain from receptor to downstream targets inside a single developmental module. *PYK10* is independently anchored as a nematode-relevant gene that accumulates in the periderm of *H. schachtii* –infected Arabidopsis roots (Schmidt, 1995) and hydrolyzes indole glucosinolates (Nakano et al., 2017), linking its NILR1-dependent induction here to the broader NILR1-sensitive indole glucosinolate module.

The *gtl1 df1* double mutant confirmed that the GTL1/DF1 module is functionally relevant to *H. schachtii* parasitism. NILR1 is expressed with highest abundance in the cortex (Shahan et al., 2022), positioning it in the cell layers most relevant for nematode approach and cortex migration (Wyss and Zunke, 1986; Siddique and Grundler, 2018). The *gtl1 df1* phenotype was not expected to match *nilr1*, and did not: NILR1 modulates only ∼3.5% of the GTL1/DF1 regulatory space (290 of 8,391 misregulated genes) in an infection-conditional, selective manner, whereas *gtl1 df1* removes the regulator constitutively across the entire network, including baseline cortex cell length (Nolan et al., 2023). A receptor that reconstituted the full *gtl1 df1* phenotype would be acting as a general regulator of cortex identity rather than a perception event tied to nematode contact. Accordingly, the two perturbations diverge: *nilr1* elevates overall susceptibility with a male-biased outcome and larger females on wild-type-sized syncytia, whereas *gtl1 df1* selectively increases female establishment (+46.9%) on smaller syncytia, with male and total counts unchanged. Why *gtl1 df1* preferentially promotes female establishment remains unclear: male- and female-associated syncytia are induced in distinct cell types and recruit surrounding cells through different patterns of hypertrophy and wall remodeling (Sobczak et al., 1997), so the altered cortical state in *gtl1 df1* may be read by invading J2s as permissive for female fate independently of subsequent syncytium development — consistent with the decoupling of female and syncytium size we observe across both mutants.

In this light, NILR1 can be viewed as a functional co-option of BRI1-clade (LRR-RLK-Xb) architecture into a pathway that contributes to pathogen resistance. Members of this receptor family are, based on comparative analyses, typically assigned to developmental roles rather than to classical immunity (Snoeck et al., 2025; Ngou et al., 2024, 2026), and they signal through a conserved brassinosteroid-type kinase cascade (Zheng et al., 2022; Ali et al., 2025). In the *H. schachtii* –Arabidopsis root system studied here, this architecture converges on the HAT7/GTL1 cortex network (Nolan et al., 2023) and is associated with reduced host permissiveness to nematode parasitism. This extends recent proposals that plant cell-surface receptors diversify beyond canonical immunity (Ngou et al., 2026) into the developmental–immune axis and raises the question of whether other receptor kinases may confer resistance through analogous developmental logic.

While our data establish a consistent picture of NILR1 as a developmentally wired receptor that feeds into a HAT7/GTL1-centred cortex module during *H. schachtii* infection (Fig. 6), several aspects of this framework remain open for exploration. We have so far focused on one Arabidopsis accession and a single *H. schachtii* population, and extending this analysis to additional host genotypes, nematode isolates and other pathosystems will show how broadly the logic uncovered here applies. Our early infection transcriptomes, based on bulk roots and discrete time points, capture robust NILR1-associated trends, but higher-resolution approaches such as cell-type-resolved or spatial transcriptomics could further refine where and when these programmes are deployed. Finally, although our genetic and network analyses support a link from NILR1 via a BR-type kinase cascade to cortex growth and wall remodeling, direct measurements of the predicted changes in cell-wall properties, growth dynamics and *in situ* ligand activity will provide valuable tests and refinements of the working model we propose.

**Figure 6:**
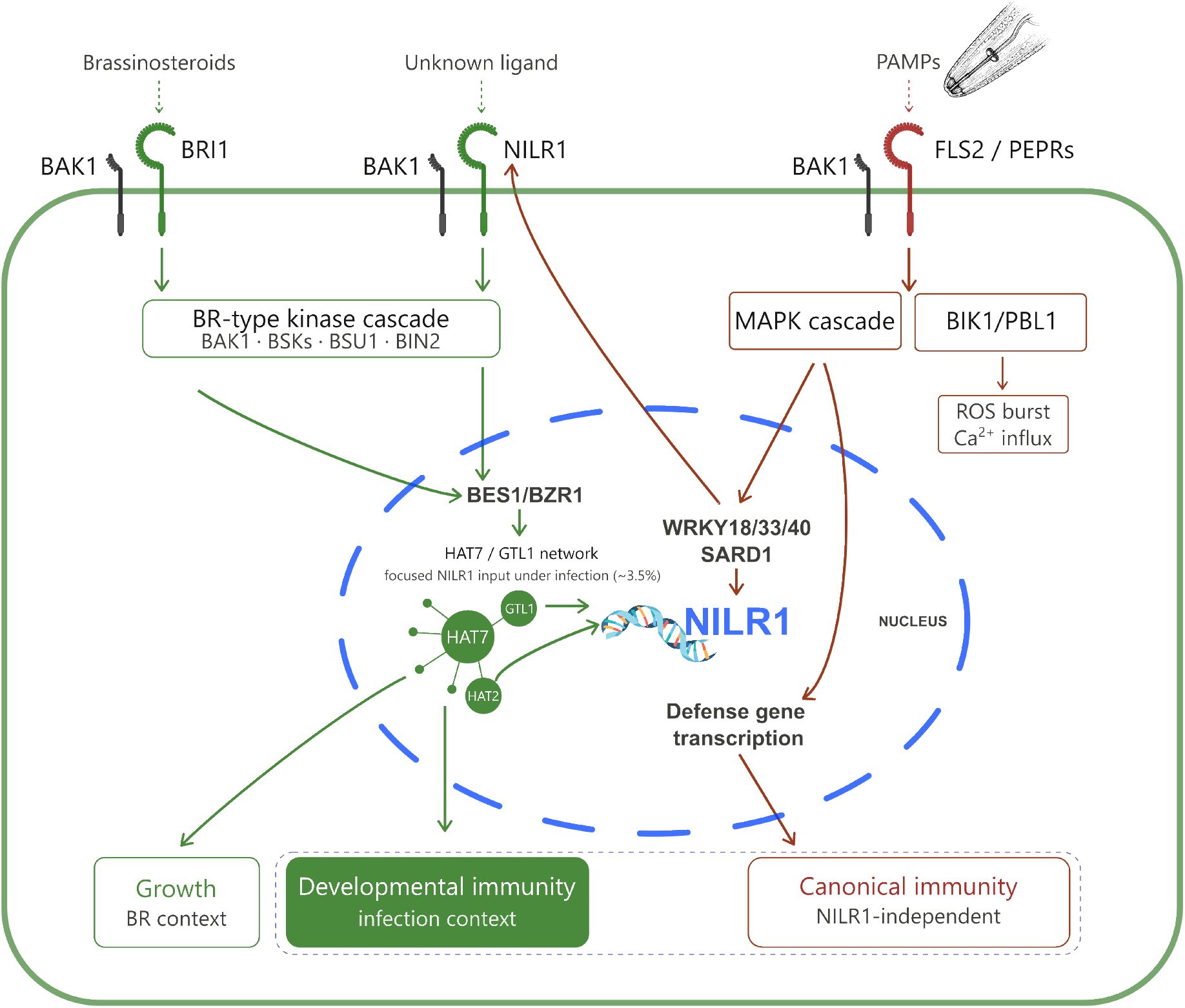
Proposed model: NILR1 is transcriptionally embedded in immunity yet routes its output through a developmental program to restrict *H. schachtii*. *Left:* in the elongating cortex, BRI1 perceives brassinosteroids and engages a BR-type kinase cascade (BAK1, BSKs, BSU1, BIN2) that activates BES1/BZR1 to drive the HAT7/GTL1 network and cortex cell elongation (Nolan et al., 2023). *Centre:* during *H. schachtii* infection, NILR1 perceives an undefined ligand and engages the same BR-type kinase cascade, providing focused transcriptional input to a subset (∼3.5%) of the HAT7/GTL1 network (developmental immunity). *Right:* PAMP perception by FLS2/PEPRs activates canonical PTI via parallel MAPK and BIK1/PBL1 branches, driving ROS and Ca^2+^ influx and, through WRKY18/33/40 and SARD1, defense gene transcription, all independently of NILR1 (canonical immunity). *Nucleus:* the *NILR1* promoter is itself a node where these programs meet: the immune regulators WRKY18/33/40 and SARD1 that drive canonical defense also drive NILR1 infection induction, while the developmental regulators GTL1 and HAT2 occupy its promoter. NILR1 is thus activated by the immune network but directs its output to the developmental module. The individual links are established, NILR1-driven BES1 dephosphorylation (Zheng et al., 2022) and HAT7/GTL1 as direct BES1/BZR1 targets (Nolan et al., 2023); their operation as a single chain during nematode infection is proposed here rather than directly demonstrated.

## Acknowledgements

We thank Trevor M. Nolan and Irene T. Liao (California Institute of Technology) for providing the *gtl1* and *gtl1 df1* mutant lines. We are grateful to Peter Marhavý for critical reading of the manuscript and helpful comments. We would further like to acknowledge Ute Schlee, Gisela Sichtermann and Josefin Mü ller for valuable technical assistance and maintenance of the nematode stock culture. Large language models (Claude, Anthropic; ChatGPT, OpenAI; Perplexity, Perplexity AI) were used to assist with drafting and editing the manuscript text and to support code development for analyses in R and Python. All scientific content, analyses, interpretations, and conclusions are the responsibility of the authors.

## Author contributions

MFE and FMWG designed the research. MFE performed and analyzed the experiments with support from SA and SN. MFE wrote the manuscript. FMWG supervised the project. All authors revised and approved the final manuscript.

## Data availability

The raw RNAseq data generated in this study has been deposited in the ENA database under Bio-Project accession PRJEB112143.

## Supporting Information

- **Supplementary Table S1.** Differentially expressed genes in wild-type (Col-0) roots at 30 min after *Heterodera schachtii* inoculation relative to mock-treated controls. This table contains the individual DEG list, including genes significantly up- or downregulated in wild-type roots at 30 min post inoculation.
- **Supplementary Table S2.** Differentially expressed genes in *nilr1* roots at 30 min after *Heterodera schachtii* inoculation relative to mock-treated controls. This table contains the individual DEG list, including genes significantly up- or downregulated in *nilr1* roots at 30 min post inoculation.
- **Supplementary Table S3.** Differentially expressed genes in wild-type (Col-0) roots at 3 h after *Heterodera schachtii* inoculation relative to mock-treated controls. This table contains the individual DEG list, including genes significantly up- or downregulated in wild-type roots at 3 h post inoculation.
- **Supplementary Table S4.** Differentially expressed genes in *nilr1* roots at 3 h after *Heterodera schachtii* inoculation relative to mock-treated controls. This table contains the individual DEG list, including genes significantly up- or downregulated in *nilr1* roots at 3 h post inoculation.
- **Supplementary Table S5.** Genotype × treatment interaction analysis at 30 min after *Heterodera schachtii* inoculation. This table lists genes ranked from the DESeq2 interaction model comparing infection responses between wild-type and *nilr1* roots at 30 min.
- **Supplementary Table S6.** Genotype × treatment interaction analysis at 3 h after *Heterodera schachtii* inoculation. This table lists genes ranked from the DESeq2 interaction model comparing infection responses between wild-type and *nilr1* roots at 3 h.
- **Supplementary Table S7.** Genotype × treatment interaction effects for a curated set of BR biosynthesis genes, core signaling components, and BR-responsive markers (*n* = 46 genes) at 30 min and 3 h post inoculation.
- **Supplementary Table S8.** List of primers used for qRT-PCR.
- **Supplementary Table S9.** GTL1 target gene sets used for gene set enrichment analysis. Predicted GTL1 (AT1G33240) target genes were extracted from Nolan et al. (2023) Supplementary Data S5 (cortex gene regulatory networks under brassinosteroid perturbation; CellOracle inference) and mapped from TAIR to Entrez Gene identifiers using org.At.tair.db. The inclusive set comprises all 294 unique GTL1 targets; the high-confidence set comprises the top 30% (88 genes) ranked by absolute mean regression coefficient. Each target is annotated with its TAIR locus, gene symbol, gene description, regression coefficient (coef mean, coef abs), edge p-value, the Nolan et al. cluster context in which the edge was inferred (BR-depleted baseline or hours after brassinolide treatment), and the cell-wall-related gene flag from the original dataset. Where a target appeared in multiple cluster contexts, one representative edge was retained per target.
- **Supplementary Table S10.** Gene set enrichment results for GTL1 high-confidence targets against the genotype × treatment interaction ranking. Rank-based GSEA was performed using clusterProfiler::GSEA with the adaptive multilevel permutation scheme implemented in fgsea, Benjamini–Hochberg p-value adjustment, and minimum and maximum gene set sizes of 10 and 5,000, respectively. Genes were ranked by the genotype × treatment interaction Wald statistic (Col-0 vs *nilr1*) at 30 min and 3 h post inoculation with *H. schachtii*. The table reports gene set size, raw enrichment score, normalized enrichment score (NES), nominal and adjusted p-values, the rank position at peak |ES|, leading-edge summary metrics (tags, list, signal), and the slash-separated Entrez identifiers of the leading-edge (core enrichment) genes.
- **Supplementary Table S11.** Leading-edge genes from the GTL1 high-confidence GSEA analysis. Leading-edge genes are members of the gene set that fall before the peak of the running enrichment score and drive the enrichment signal. Genes are listed for both timepoints (33 at 30 min; 38 at 3 h), annotated with TAIR locus, gene symbol, and gene description, and reported with their genotype × treatment interaction Wald statistic and rank position in the full ranked gene list. Genes are sorted by interaction statistic in ascending order, so the most strongly NILR1-dependent genes appear first.
- **Supplementary Table S12.** Baseline genotype effect (*nilr1* vs WT) under mock conditions.
- **Supplementary Table S13**. Statistical analysis of the Col-0 vs *nilr1* infection phenotype (Fig. 1). Full results of the negative-binomial models for nematode counts (females, males, total) and the log-linear models for female and syncytium size, each with genotype as a fixed effect and biological replicate as a fixed block (*n* = 3 biological replicates). For every panel the table reports the *nilr1*/Col-0 effect size (incidence rate ratio for counts, geometric-mean ratio for size) with 95% confidence interval and *P* value, the genotype × replicate interaction (consistency of the effect across replicates), and a conservative replicate-level paired *t* -test on per-replicate genotype means. Per-group descriptive statistics are included.
- **Supplementary Table S14**. Statistical analysis of the multi-genotype (GTL1/DF1) infection phenotype (Fig. 5). Full results for the Col-0, *nilr1*, *gtl1* and *gtl1 df1* comparison (*n* = 4 biological replicates). Female and male counts were analyzed jointly by a negative-binomial GLMM (Genotype × Sex, with plant nested within replicate); total counts by an analogous Genotype-only model. Sheets give the Type II Wald *χ*^2^ tests, estimated marginal means, Tukey-adjusted pairwise comparisons, and S^̌^ idák-adjusted compact letter displays. Female and syncytium sizes are reported with two models: the primary linear mixed model on individual log-transformed measurements (Option B) and a conservative Tukey test on replicate-level means (Option A). A key-results sheet summarises each genotype-versus-Col-0 contrast as a percentage change with its *P* value.
- **Supplementary Table S15.** NILR1-dependent transcription factors at 30 min (*n* = 14) and 3 h (*n* = 16) post inoculation and their mapping onto the brassinosteroid-responsive cortex elongation GRN of Nolan et al., 2023. For each TF, the table reports timepoint, Entrez ID, TAIR locus, gene symbol, and—where present in the network—cluster, node role, degree, and centrality measures.
- **Supplementary Table S16.** Functional categorisation of the 290 NILR1-dependent candidate genes (interaction shrunken log_2_FC ≤ −0.2) at 3 h post inoculation, with per-gene shrunken and raw log_2_ fold changes, adjusted *p*-value, baseMean, TF family, and functional annotation.

**Figure S1:**
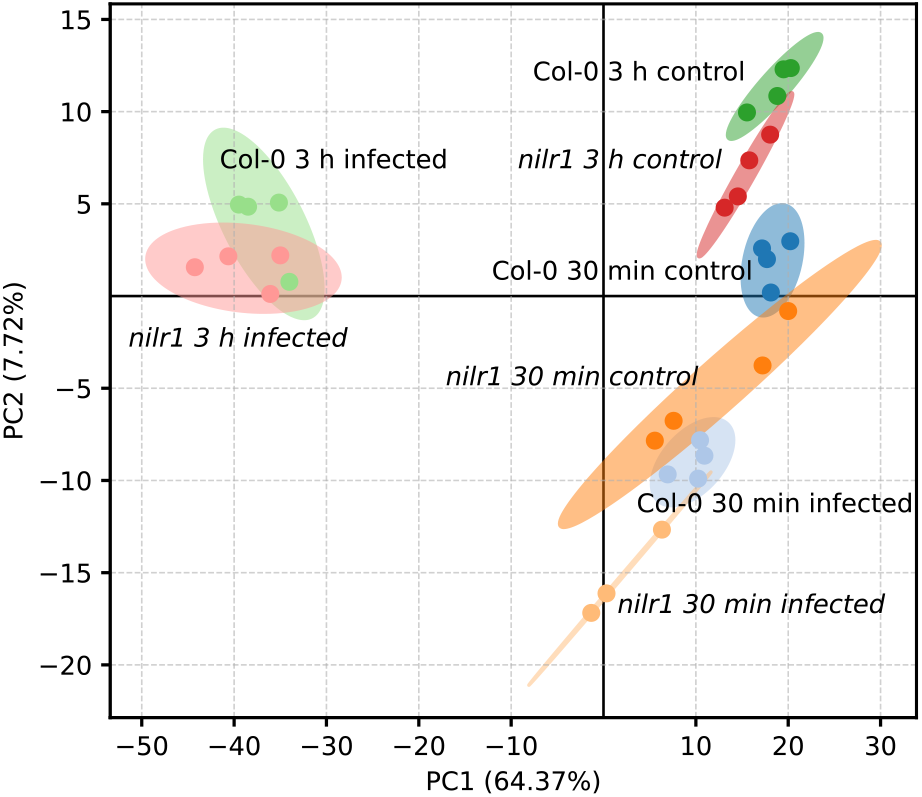
Principal component analysis of transcriptomes from infected and mock-treated roots. PC1 (64.37%) primarily reflects nematode-induced transcriptional reprogramming, whereas PC2 (7.72%) captures temporal effects. Ellipses indicate 95% confidence intervals.

**Figure S2:**
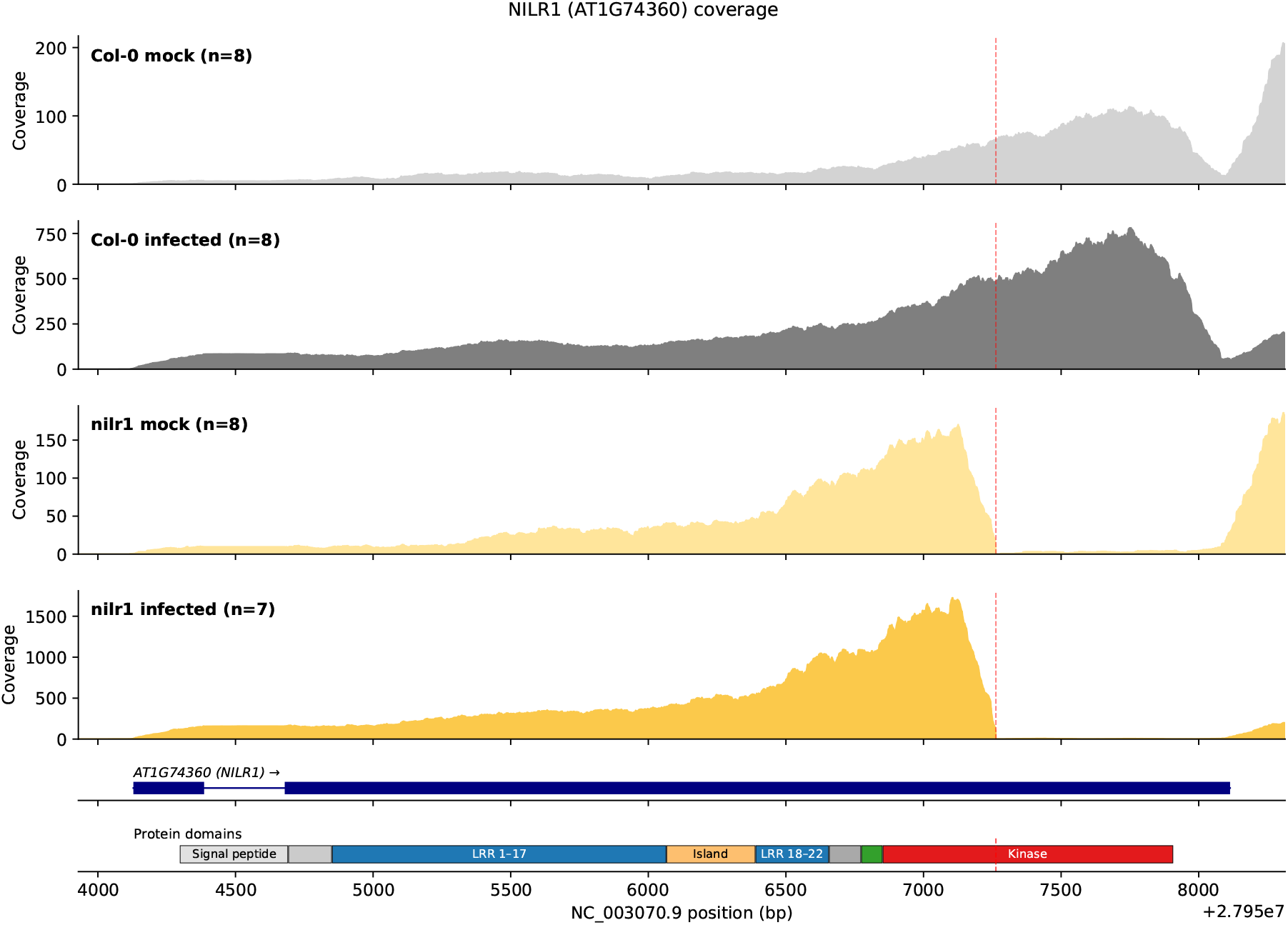
RNAseq coverage of NILR1. RNA-seq read coverage across the *NILR1* locus (*AT1G74360*, T-DNA insertion line SAIL 865 E07), averaged within each genotype × treatment combination (timepoints 30 min and 3 h pooled; n = 8 for Col-0 mock, Col-0 infected, and *nilr1* mock; n = 7 for *nilr1* infected). Wild-type (Col-0) samples show continuous coverage across the 3,986-bp gene body. *nilr1* samples show a sharp coverage dropout at chr1:27,957,263 (red dashed line), corresponding to amino acid 891 of the 1,106-aa NILR1 receptor within the Ser/Thr kinase domain (Mendy et al., 2017). Gene model (navy) and protein domain architecture (Mendy et al., 2017) are shown below. Coverage y-axes are scaled independently per panel.

**Figure S3:**
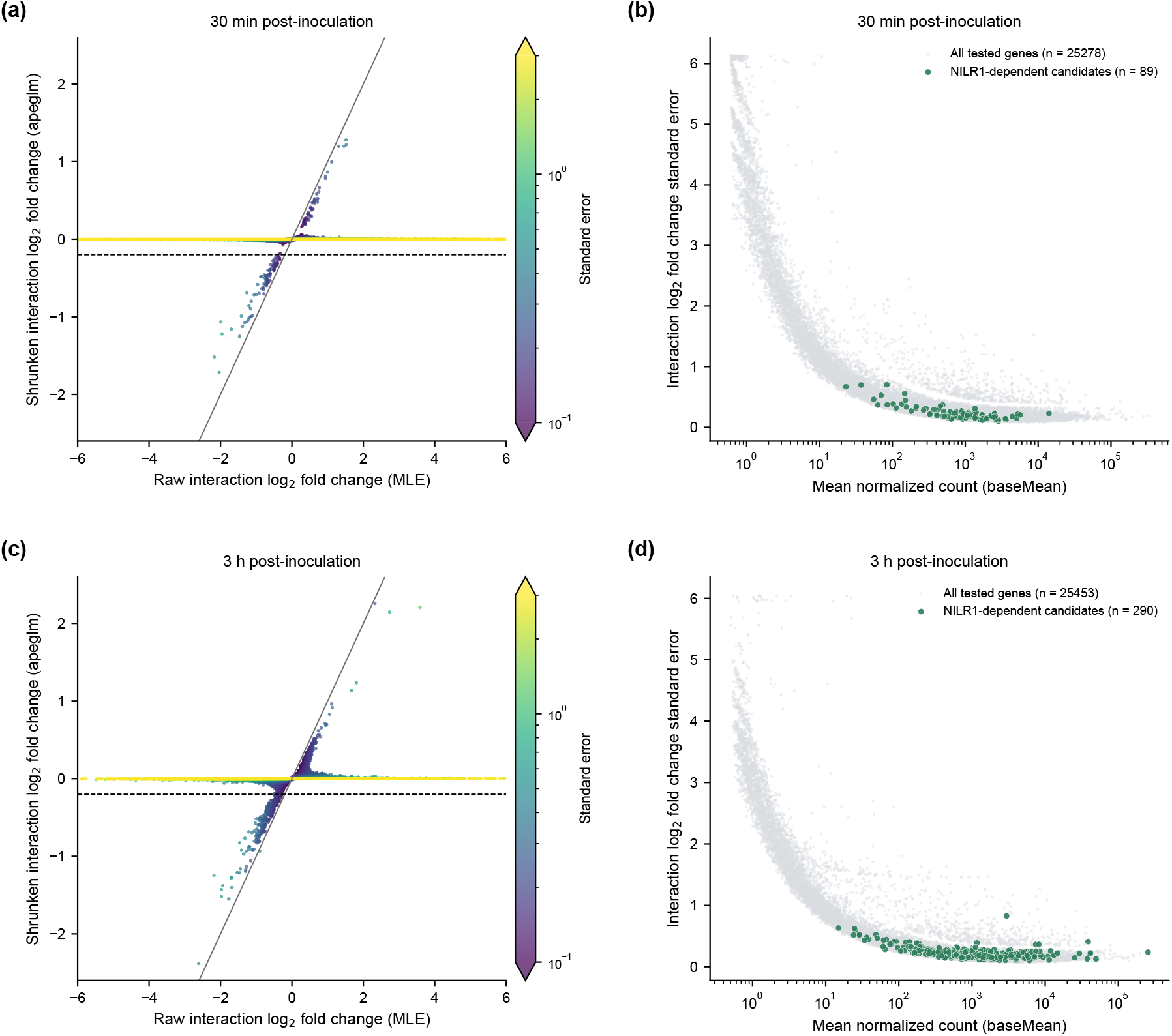
Effect-size shrinkage of the genotype × treatment interaction estimates is governed by per-gene uncertainty. Genotype × treatment interaction effects were estimated with DESeq2 (model ∼ genotype + treatment + genotype:treatment; *nilr1* vs wild type) and stabilised using the apeglm shrinkage estimator (Zhu et al., 2019), separately at each post-inoculation timepoint. NILR1-dependent candidates were defined as genes with shrunken interaction log_2_ fold change ≤ −0.2. **(a, c)** Raw maximum-likelihood estimate (MLE) of the interaction log_2_ fold change (x-axis) versus the apeglm-shrunken estimate (y-axis) for all tested genes at 30 min (a) and 3 h (c) post-inoculation, coloured by the standard error of the interaction estimate. The solid line denotes the identity (no shrinkage); the horizontal dashed line marks the candidate threshold (−0.2). Estimates with large standard errors are collapsed toward zero, whereas estimates with small standard errors are retained close to the identity line, demonstrating that shrinkage is determined by estimate precision rather than by raw effect magnitude. **(b, d)** Standard error of the interaction log_2_ fold change versus mean normalised count (baseMean) at 30 min (b) and 3 h (d). NILR1-dependent candidates (green) are distributed across the high-expression, low-uncertainty region of the dataset (all candidates have baseMean ≥ 20; median standard error 0.21 at both timepoints), confirming that the candidate set comprises adequately sampled, precisely estimated genes.

**Figure S4:**
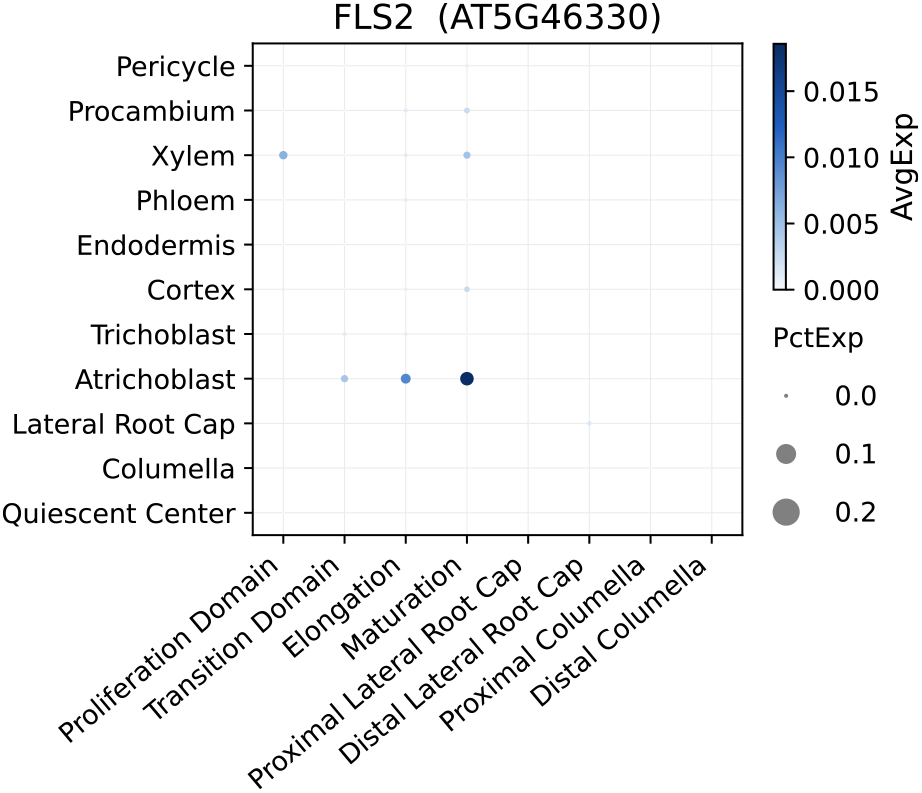
Cell-type-resolved expression of *FLS2* (AT5G46330) across root cell types, based on published single-cell transcriptomic data (Shahan et al., 2022). *FLS2* displays lower overall transcript abundance under non-elicited conditions than NILR1, with expression enriched in atrichoblasts and vascular-associated cell types.

**Figure S5:**
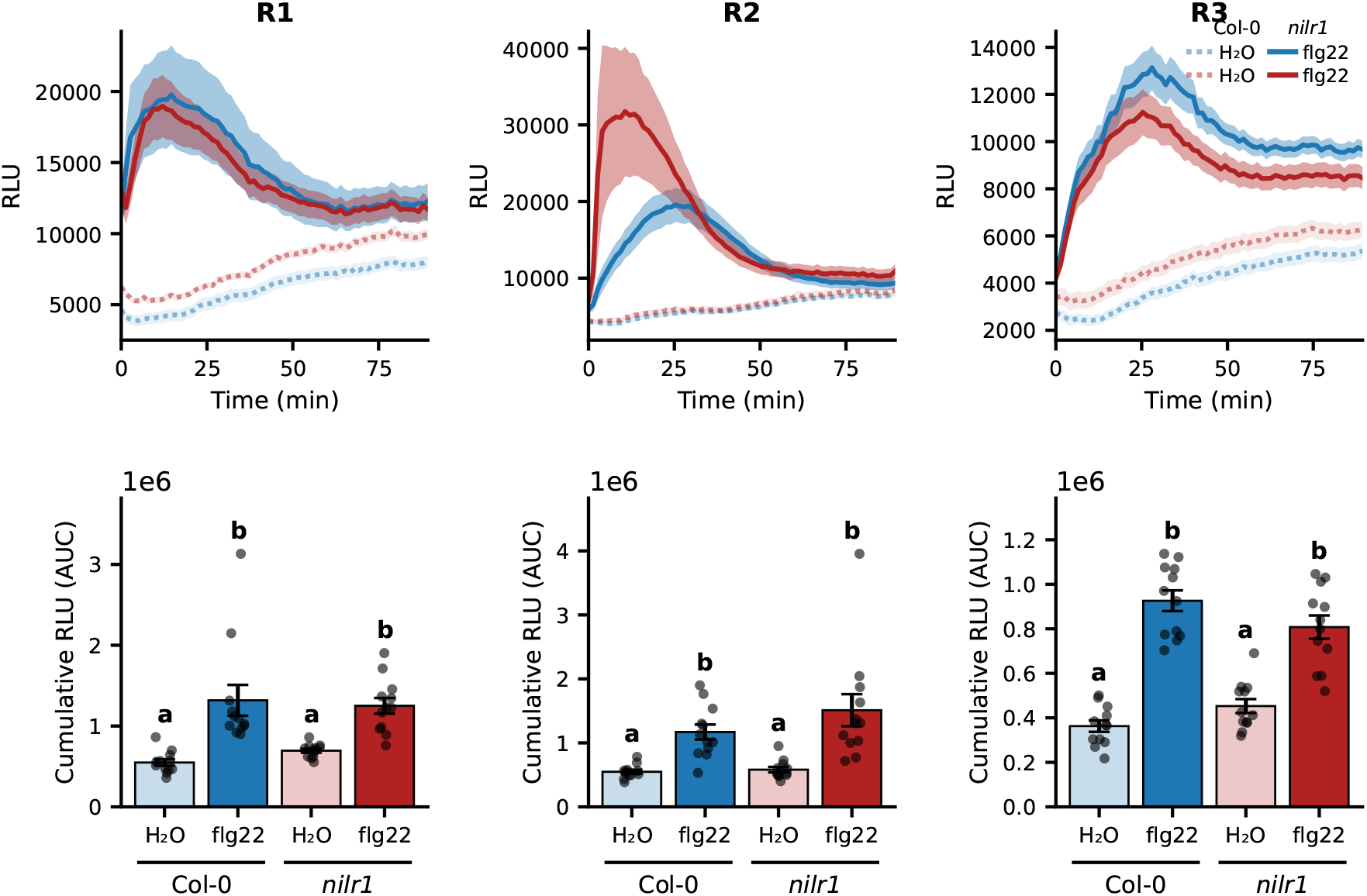
Apoplastic ROS production in Col-0 and *nilr1* leaf discs across three independent biological replicates. Leaf discs from 12-day-old Col-0 and *nilr1* seedlings were treated with 0.5 µM flg22 or water, and luminol-based chemiluminescence was recorded at 1 min 20 sec intervals for 90 min using a TECAN Infinite 200 Pro microplate reader. **(Top row)** ROS production kinetics for replicates R1–R3. Traces show the mean of *n* = 12 leaf discs per treatment; shaded areas indicate ± SEM. **(Bottom row)** Cumulative ROS production over 90 min, calculated as the area under the curve (AUC) of the kinetic traces shown above. Bars represent mean ± SEM; dots show individual leaf discs (*n* = 12). Statistical analysis was performed by two-way ANOVA with genotype and treatment as fixed factors, followed by Tukey’s HSD post-hoc test. Different letters above bars indicate statistically significant differences between groups within each replicate (*p <* 0.05). In all three replicates, the genotype × treatment interaction was not significant, and flg22-treated samples did not differ between Col-0 and *nilr1*, confirming that NILR1 is dispensable for flg22-triggered ROS production.

**Figure S6:**
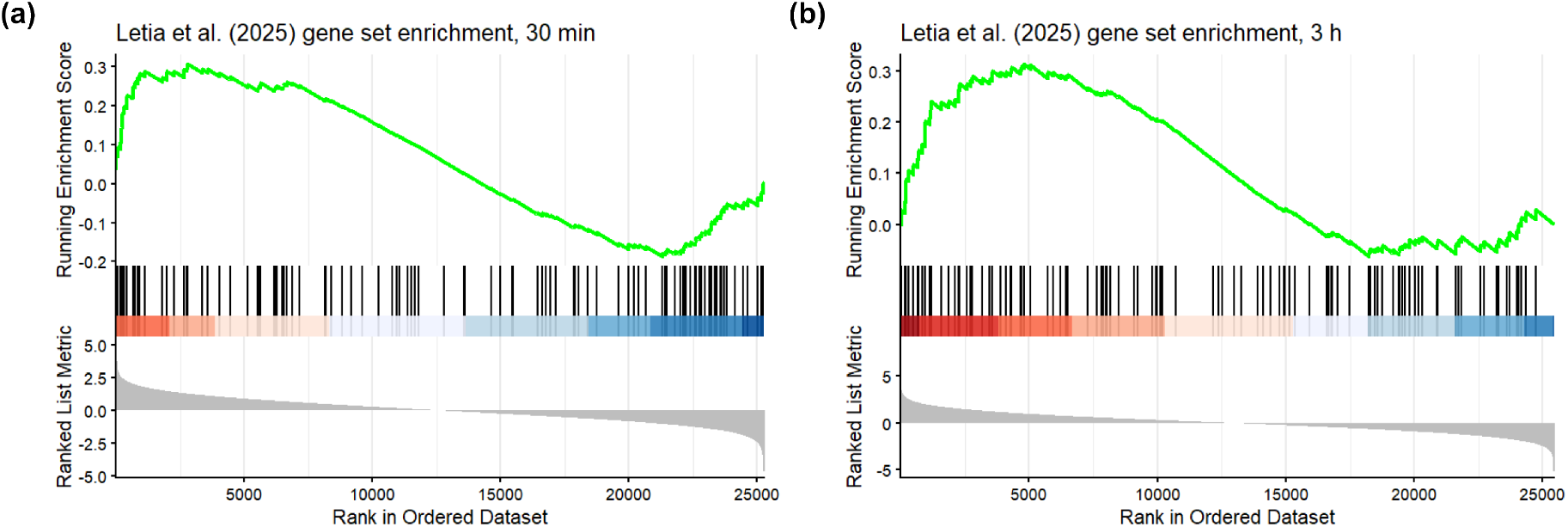
Ascaroside-responsive genes align with NILR1-independent components of the nematode response. Gene set enrichment analysis of the Letia et al. (2025) ascaroside-responsive gene set against the genotype × treatment interaction ranking at **(a)** 30 min (NES = +1.42, *p* = 0.022) and **(b)** 3 h (NES = +1.48, adjusted *p* = 0.010) post inoculation with *H. schachtii*. Genes were ranked by the DESeq2 interaction Wald statistic (Col-0 vs *nilr1*); positive statistics indicate enhanced or de-repressed responses in *nilr1*, negative statistics indicate attenuated responses. Enrichment of ascaroside-responsive genes toward positive interaction statistics at both time points indicates preferential alignment with NILR1-independent components of the infection response.

**Figure S7:**
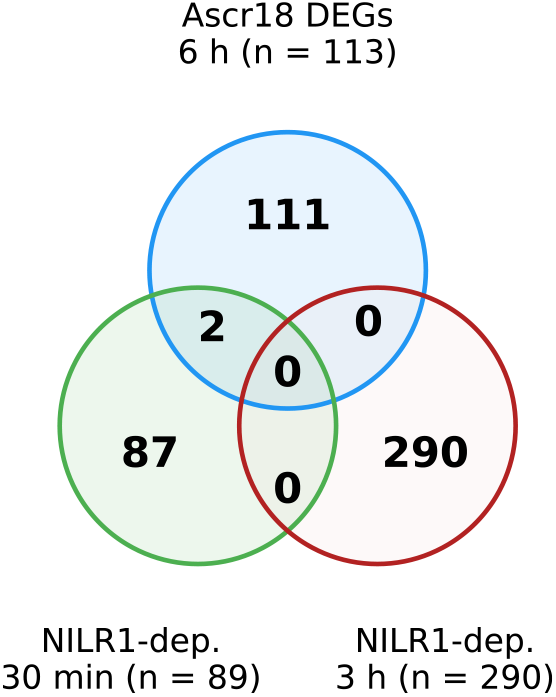
Overlap between ascaroside-responsive DEGs (Letia et al., 2025; blue, *n* = 113) and NILR1-dependent candidate genes at 30 min (green, *n* = 89) and 3 h (red, *n* = 290) post inoculation. Numbers indicate gene counts in each set and their intersections. Gene-level overlap was minimal — only *HAT2* and *AT3G59320* were shared at 30 min, and no genes overlapped at 3 h — although the two programs converge on shared functional categories including cell wall polysaccharide metabolism (Letia et al., 2025).

**Figure S8:**
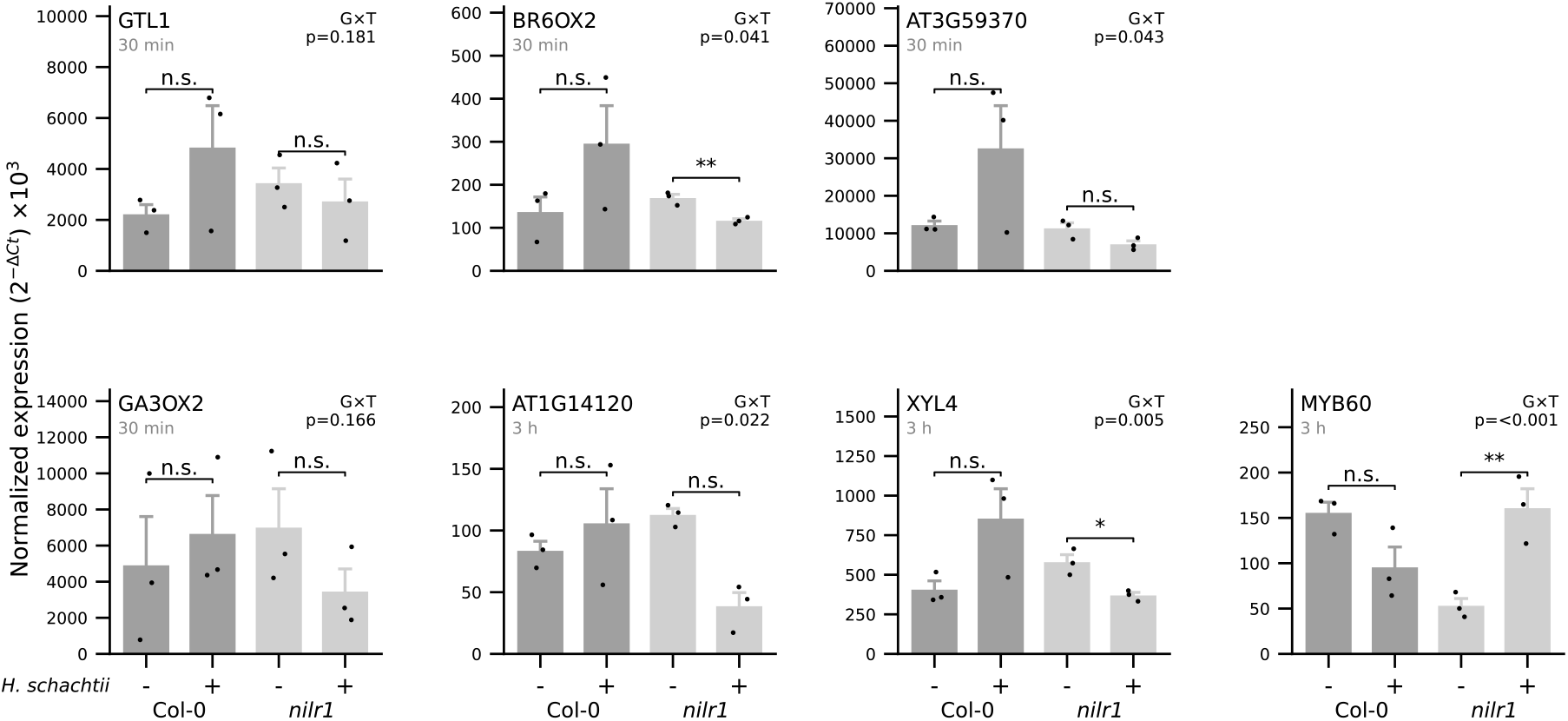
Additional qRT-PCR validation of NILR1-dependent candidate genes. Expression of NILR1-dependent candidate genes in wild-type (Col-0) and *nilr1* roots upon inoculation with *H. schachtii*, quantified at the timepoint indicated within each panel (30 min or 3 h post inoculation). These genes were identified as NILR1-dependent candidates in the genotype × treatment interaction analysis. Expression patterns by qRT-PCR are consistent in direction with the RNA-seq results, although effect sizes for individual within-genotype comparisons are variable, reflecting the limited power of *n* = 3. Dark grey bars, Col-0; light grey bars, *nilr1*. Within-genotype comparisons (control vs infected) were assessed using Welch’s two-sided *t*-test on Δ*Ct* values; significance is indicated above each bracket (* *P <* 0.05, ** *P <* 0.01, *** *P <* 0.001, n.s. not significant). G×T *P* - values indicate the significance of the genotype × treatment interaction (two-factor linear model on Δ*Ct*). Bars represent the mean of three biological replicates (*n* = 3) ± SE; individual data points are overlaid.

**Figure S9:**
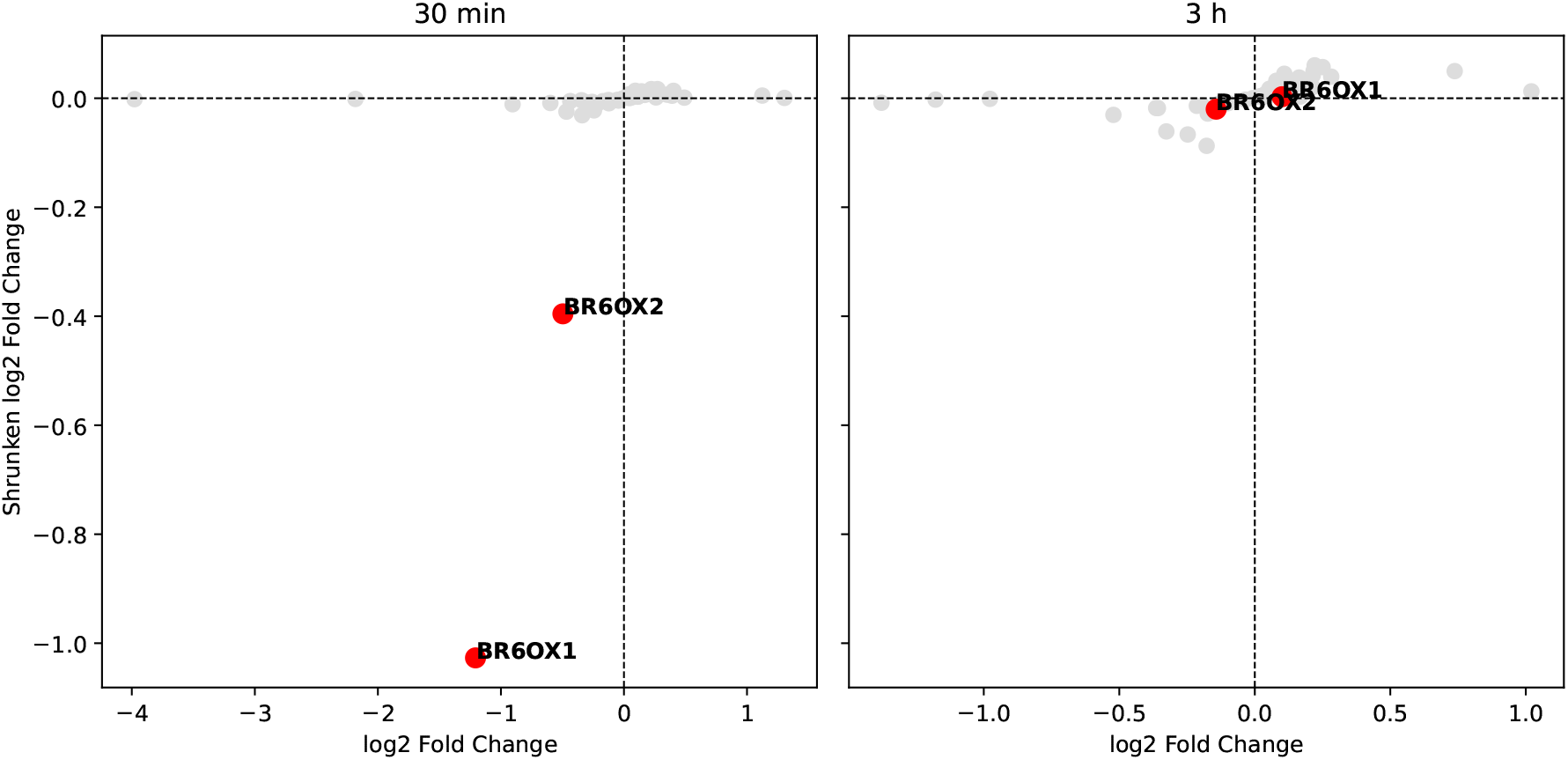
NILR1-dependent effects on brassinosteroid pathway genes during early nematode infection. Scatterplots show interaction log2 fold changes (x-axis) versus shrunken log2 fold changes (y-axis) for a curated set of brassinosteroid (BR) biosynthesis genes, core BR signaling components, and BR-responsive transcriptional regulators at 30 min (left) and 3 h (right) after *Heterodera schachtii* inoculation. Each point represents one gene. Red points highlight the late BR biosynthetic enzymes *BR6OX1* (*CYP85A1*) and *BR6OX2* (*CYP85A2*). At 30 min post inoculation, both genes show pronounced negative interaction effects, whereas interaction effects for the remaining BR-related genes cluster near zero. At 3 h post inoculation, interaction effects across the BR gene set are minimal, indicating the absence of sustained NILR1-dependent transcriptional responses in the BR pathway. Dashed lines indicate zero log2 fold change.

**Figure S10:**
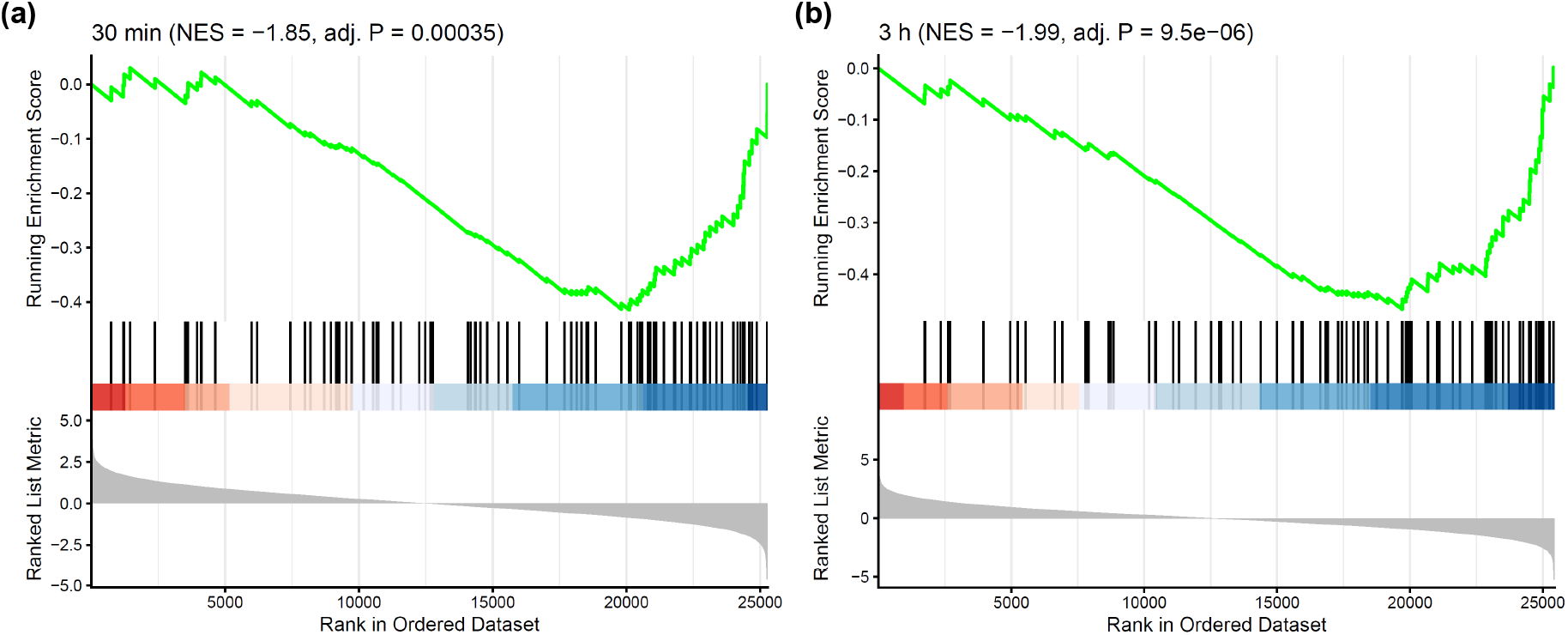
Enrichment of high-confidence GTL1 target genes among NILR1-dependent transcriptional responses at 30 min post inoculation. Gene set enrichment analysis of high-confidence GTL1 target genes (top 30% of GTL1 targets in the Nolan et al. (2023) cortex gene regulatory network, *n* = 88) against the genotype × treatment interaction ranking (Col-0 vs *nilr1*) at 30 min post inoculation with *H. schachtii*. Genes were ranked by the DESeq2 interaction Wald statistic; negative statistics indicate attenuated infection responses in *nilr1*. The running enrichment score (green) trends negative, indicating that GTL1 targets are concentrated among NILR1-dependent genes (NES = −1.85, adjusted *P* = 3.5 × 10*^−^*^4^; 33 leading-edge genes). **(b)** Enrichment of GTL1 targets toward negative interaction statistics is more pronounced than at 30 min (NES = −1.99, adjusted *P* = 9.5 × 10*^−^*^6^; 38 leading-edge genes), consistent with progressive engagement of the GTL1 module during early infection. The lower panel shows the distribution of interaction statistics across the dataset.

## References

1. Ali, K., Li, W., & Wu, G. (2025). Kinase domain diversification drives specificity in BRI1 and non-BRI1 RLKs in brassinosteroid signaling. Plant Science, 357 (February), 112531. 10.1016/j.plantsci.2025.112531

2. Birkenbihl, R. P., Kracher, B., Roccaro, M., & Somssich, I. E. (2017). Induced Genome-Wide Binding of Three Arabidopsis WRKY Transcription Factors during Early MAMP-Triggered Immunity. The Plant Cell, 29(1), 20–38. 10.1105/tpc.16.00681

3. Bjornson, M., Pimprikar, P., Nürnberger, T., & Zipfel, C. (2021). The transcriptional landscape of *Arabidopsis thaliana* pattern-triggered immunity. Nature Plants, 7 (May). 10.1038/s41477-021-00874-5

4. Breuer, C., Kawamura, A., Ichikawa, T., Tominaga-Wada, R., Wada, T., Kondou, Y., Muto, S., Matsui, M., & Sugimoto, K. (2009). The trihelix transcription factor GTL1 regulates ploidy-dependent cell growth in the *Arabidopsis* trichome. Plant Cell, 21(8), 2307–2322. 10.1105/tpc.109.068387

5. Chen, Y., Zheng, X., Li, J., & Li, X. (2025). Regulation of cadmium accumulation in plants by the transcription factor GTL1: Potential for minimizing grain cadmium. Journal of Hazardous Materials, 499(August), 140302. 10.1016/j.jhazmat.2025.140302

6. Chinchilla, D., Bauer, Z., Regenass, M., Boller, T., & Felix, G. (2006). The *Arabidopsis* receptor kinase FLS2 binds flg22 and determines the specificity of flagellin perception. Plant Cell, 18(2), 465– 476. 10.1105/tpc.105.036574

7. Davidsson, P., Broberg, M., Kariola, T., Sipari, N., Pirhonen, M., & Palva, E. T. (2017). Short oligogalacturonides induce pathogen resistance-associated gene expression in Arabidopsis thaliana. BMC Plant Biology, 17 (1), 19. 10.1186/s12870-016-0959-1

8. Felix, G., Duran, J. D., Volko, S., & Boller, T. (1999). Plants have a sensitive perception system for the most conserved domain of bacterial flagellin. Plant Journal, 18(3), 265–276. 10.1046/j.1365-313X.1999.00265.x

9. Gombos, M., Raynaud, C., Nomoto, Y., Molnár, E., Brik-Chaouche, R., Takatsuka, H., Zaki, A., Bernula, D., Latrasse, D., Mineta, K., Nagy, F., He, X., Iwakawa, H., Őszi, E., An, J., Suzuki, T., Papdi, C., Bergis, C., Benhamed, M., Bögre, L., Ito, M., & Magyar, Z. (2023). The canonical E2Fs together with RETINOBLASTOMA-RELATED are required to establish quiescence during plant development. Communications Biology, 6(1), 1–15. 10.1038/s42003-023-05259-2

10. Huang, L., Yuan, Y., Lewis, C., Kud, J., Kuhl, J. C., Caplan, A., Dandurand, L. M., Zasada, I., & Xiao, F. (2023). NILR1 perceives a nematode ascaroside triggering immune signaling and resistance. Current Biology, 33(18), 3992–3997.e3. 10.1016/j.cub.2023.08.017

11. Huang, L., Yuan, Y., Ramirez, C., Zhao, Z., Chen, L., Griebel, T., Kud, J., Kuhl, J. C., Caplan, A., Dandurand, L.-M., & Xiao, F. (2024). A receptor for dual ligands governs plant immunity and hormone response and is targeted by a nematode effector. Proceedings of the National Academy of Sciences, 121(42). 10.1073/pnas.2412016121

12. Iino, E., Kadota, Y., Maki, N., Sato, K., Ono, E., Ishihama, N., Rugen-Hankey, M., Wei, S., Ngou, B. P. M., Kumakura, N., Schmid, M. W., Eves-Van den Akker, S., Suzuki, T., Uehara, T., & Shirasu, K. (2025). A trehalase-derived MAMP triggers LecRK-V–mediated immune responses in *Arabidopsis*. Science Advances, 11(31), 1–14. 10.1126/sciadv.adv8896

13. Jin, J., Tian, F., Yang, D.-C., Meng, Y.-Q., Kong, L., Luo, J., & Gao, G. (2017). PlantTFDB 4.0: toward a central hub for transcription factors and regulatory interactions in plants. Nucleic Acids Research, 45(D1), D1040–D1045. 10.1093/nar/gkw982

14. Jones, J. D., & Dangl, J. L. (2006). The plant immune system. Nature, 444(7117), 323–329. 10.1038/nature05286

15. Jones, J. D., Staskawicz, B. J., & Dangl, J. L. (2024). The plant immune system: From discovery to deployment. Cell, 187 (9), 2095–2116. 10.1016/j.cell.2024.03.045

16. Kim, D., Paggi, J. M., Park, C., Bennett, C., & Salzberg, S. L. (2019). Graph-based genome alignment and genotyping with HISAT2 and HISAT-genotype. Nature Biotechnology, 37 (8), 907–915. 10.1038/s41587-019-0201-4

17. Lamers, J., Zhang, Y., van Zelm, E., Leong, C. K., Meyer, A. J., de Zeeuw, T., Verstappen, F., Veen, M., Deolu-Ajayi, A. O., Gommers, C. M. M., & Testerink, C. (2025). Abscisic acid signaling gates salt-induced responses of plant roots. Proceedings of the National Academy of Sciences, 122(6), 2017. 10.1073/pnas.2406373122

18. Letia, S., Bhattacharyya, S., Mendy, B., Vothknecht, U. C., von Reuss, S. H., Inada, M., Grundler, F. M. W., & Hasan, M. S. (2025). Ascaroside#18 Promotes Plant Defence by Repressing Auxin Signalling. Physiologia Plantarum, 177 (4). 10.1111/ppl.70386

19. Liao, Y., Smyth, G. K., & Shi, W. (2014). FeatureCounts: An efficient general purpose program for assigning sequence reads to genomic features. Bioinformatics, 30(7), 923–930. 10.1093/bioinformatics/btt656

20. Love, M. I., Huber, W., & Anders, S. (2014). Moderated estimation of fold change and dispersion for RNA-seq data with DESeq2. Genome Biology, 15(12), 1–21. 10.1186/s13059-014-0550-8

21. Mano, N. A., Shaikh, M. A., Widhalm, J. R., Yoo, C. Y., & Mickelbart, M. V. (2024). Transcriptional repression of *GTL1* under water-deficit stress promotes anthocyanin biosynthesis to enhance drought tolerance. Plant Direct, 8(5). 10.1002/pld3.594

22. Manohar, M., Sistenich, A., Liu, S., Baby, S., Chen, S., Dejonghe, W., Kumari, A., Luna, E., Levecque, S., Manosalva, P. M., Leach, J. E., Wang, X., Kachroo, A., Kogel, K.-H., Conrath, U., Schroeder, F. C., & Klessig, D. F. (2026). The conserved nematode pheromone ascr#18 primes plant immunity. Communications Biology. 10.1038/s42003-026-10211-1

23. Matsushima, R., Fukao, Y., Nishimura, M., & Hara-Nishimura, I. (2004). NAI1 Gene Encodes a Basic-Helix-Loop-Helix–Type Putative Transcription Factor That Regulates the Formation of an Endoplasmic Reticulum–Derived Structure, the ER Body. The Plant Cell, 16(6), 1536–1549. 10.1105/tpc.021154

24. Mendy, B., Wang’ombe, M. W., Radakovic, Z. S., Holbein, J., Ilyas, M., Chopra, D., Holton, N., Zipfel, C., Grundler, F. M., & Siddique, S. (2017). *Arabidopsis* leucine-rich repeat receptor–like kinase NILR1 is required for induction of innate immunity to parasitic nematodes. PLoS Pathogens, 13(4), 1–22. 10.1371/journal.ppat.1006284

25. Miya, A., Albert, P., Shinya, T., Desaki, Y., Ichimura, K., Shirasu, K., Narusaka, Y., Kawakami, N., Kaku, H., & Shibuya, N. (2007). CERK1, a LysM receptor kinase, is essential for chitin elicitor signaling in *Arabidopsis*. Proceedings of the National Academy of Sciences of the United States of America, 104(49), 19613–19618. 10.1073/pnas.0705147104

26. Nakano, R. T., Piślewska-Bednarek, M., Yamada, K., Edger, P. P., Miyahara, M., Kondo, M., Böttcher, C., Mori, M., Nishimura, M., Schulze-Lefert, P., Hara-Nishimura, I., & Bednarek, P. (2017). PYK10 myrosinase reveals a functional coordination between endoplasmic reticulum bodies and glucosinolates in *Arabidopsis thaliana*. The Plant Journal, 89(2), 204–220. 10.1111/tpj.13377

27. Ngou, B. P. M., Kadota, Y., & Shirasu, K. (2026). Plant cell surface receptors. The Plant Journal, 125(6), 1–24. 10.1111/tpj.70800

28. Ngou, B. P. M., Wyler, M., Schmid, M. W., Kadota, Y., & Shirasu, K. (2024). Evolutionary trajectory of pattern recognition receptors in plants. Nature Communications, 15(1), 308. 10.1038/s41467-023-44408-3

29. Nolan, T. M., Vukašinović, N., Hsu, C. W., Zhang, J., Vanhoutte, I., Shahan, R., Taylor, I. W., Green-street, L., Heitz, M., Afanassiev, A., Wang, P., Szekely, P., Brosnan, A., Yin, Y., Schiebinger, G., Ohler, U., Russinova, E., & Benfey, P. N. (2023). Brassinosteroid gene regulatory networks at cellular resolution in the *Arabidopsis* root. Science, 379(6639). 10.1126/science.adf4721

30. Nolan, T. M., Vukašinović, N., Liu, D., Russinova, E., & Yin, Y. (2020). Brassinosteroids: Multidimensional Regulators of Plant Growth, Development, and Stress Responses. The Plant Cell, 32(2), 295–318. 10.1105/tpc.19.00335

31. Oh, E., Zhu, J.-Y., Bai, M.-Y., Arenhart, R. A., Sun, Y., & Wang, Z.-Y. (2014). Cell elongation is regulated through a central circuit of interacting transcription factors in the *Arabidopsis* hypocotyl. eLife, 3(3), 1–19. 10.7554/eLife.03031

32. O’Malley, R. C., Huang, S.-s. C., Song, L., Lewsey, M. G., Bartlett, A., Nery, J. R., Galli, M., Gallavotti, A., & Ecker, J. R. (2016). Cistrome and Epicistrome Features Shape the Regulatory DNA Landscape. Cell, 165(5), 1280–1292. 10.1016/j.cell.2016.04.038

33. Pijnacker, A., Wang, Y., Willig, J.-j., Mars, J., Werner, S., Adema, K., Smant, G., Korswagen, H. C., & Lozano-Torres, J. L. (2026). RNA tomography reveals spatial gene expression maps of *Arabidopsis thaliana* roots infected with *Heterodera schachtii*. New Phytologist, 249(1), 588– 602. 10.1111/nph.70674

34. Ramirez, V. E., Shuai, H., Hwu, F.-y., Hazarika, R. R., Tao, C.-n., Choi, S., Piecyk, R. S., Wudy, S. I., Gigl, M., Bagnoli, J. W., Brajkovic, S., Albertos, P., Liang, Y., Keymer, A., Dawid, C., Enard, W., Vlot, A. C., Gutjahr, C., Parniske, M., Kuster, B., Sieberer, T., Ludwig, C., Zipfel, C., Ton, J., Johannes, F., & Poppenberger, B. (2026). Brassinosteroid-regulated transcription factors confer epigenetic changes that repress plant immunity. bioRxiv. 10.64898/2026.04.09.717176

35. Rymen, B., Kawamura, A., Lambolez, A., Inagaki, S., Takebayashi, A., Iwase, A., Sakamoto, Y., Sako, K., Favero, D. S., Ikeuchi, M., Suzuki, T., Seki, M., Kakutani, T., Roudier, F., & Sugimoto, K. (2019). Histone acetylation orchestrates wound-induced transcriptional activation and cellular reprogramming in Arabidopsis. Communications Biology, 2(1), 404. 10.1038/s42003-019-0646-5

36. Safaeizadeh, M., Boller, T., & Becker, C. (2024). Comparative RNA-seq analysis of *Arabidopsis thaliana* response to *At* Pep1 and flg22, reveals the identification of PP2-B13 and ACLP1 as new members in pattern-triggered immunity. PLoS One, 19(6 June), 1–33. 10.1371/journal.pone.0297124

37. Saura-Sanchez, M., Gómez Rojas, A., Deveux, M., Minne, M., Grones, C., Eekhout, T., Abril-Urias, P., Van Bel, M., Tenorio Berrio, R., Vandepoele, K., Escobar Lucas, C., Beeckman, T., De Rybel, B., & Kyndt, T. (2025). Conserved transcriptional reprogramming in nematode infected root cells. bioRxiv. 10.1101/2025.09.05.674408

38. Savary, S., Willocquet, L., Pethybridge, S. J., Esker, P., McRoberts, N., & Nelson, A. (2019). The global burden of pathogens and pests on major food crops. Nature Ecology and Evolution, 3(3), 430–439. 10.1038/s41559-018-0793-y

39. Schmidt, K.-P. (1995). Proteinanalytische Charakterisierung pathogenesespezifischer Vorgä nge im Wurzelgewebe von Arabidopsis thaliana nach Infektion mit dem Rü benzystennematoden Heterodera schachtii [Doctoral dissertation, Christian-Albrechts-Universitä t zu Kiel].

40. Shahan, R., Hsu, C. W., Nolan, T. M., Cole, B. J., Taylor, I. W., Greenstreet, L., Zhang, S., Afanassiev, A., Vlot, A. H. C., Schiebinger, G., Benfey, P. N., & Ohler, U. (2022). A single-cell *Arabidopsis* root atlas reveals developmental trajectories in wild-type and cell identity mutants. Developmental Cell, 57 (4), 543–560.e9. 10.1016/j.devcel.2022.01.008

41. Shibata, M., Breuer, C., Kawamura, A., Clark, N. M., Rymen, B., Braidwood, L., Morohashi, K., Busch, W., Benfey, P. N., Sozzani, R., & Sugimoto, K. (2018). GTL1 and DF1 regulate root hair growth through transcriptional repression of *ROOT HAIR DEFECTIVE 6-LIKE 4* in *Arabidopsis*. Development (Cambridge), 145(3). 10.1242/dev.159707

42. Shibata, M., Favero, D. S., Takebayashi, R., Takebayashi, A., Kawamura, A., Rymen, B., Hosokawa, Y., & Sugimoto, K. (2022). Trihelix transcription factors GTL1 and DF1 prevent aberrant root hair formation in an excess nutrient condition. New Phytologist, 235(4), 1426–1441. 10.1111/nph.18255

43. Siddique, S., Coomer, A., Baum, T., & Williamson, V. M. (2022). Recognition and Response in Plant–Nematode Interactions. Annual Review of Phytopathology, 60(1), 1–20. 10.1146/annurev-phyto-020620-102355

44. Siddique, S., & Grundler, F. M. (2018). Parasitic nematodes manipulate plant development to establish feeding sites. Current Opinion in Microbiology, 46, 102–108. 10.1016/j.mib.2018.09.004

45. Snoeck, S., Johanndrees, O., Nürnberger, T., & Zipfel, C. (2025). Plant pattern recognition receptors: from evolutionary insight to engineering. Nature Reviews Genetics, 26(4), 268–278. 10.1038/s41576-024-00793-z

46. Sobczak, M., Golinowski, W., & Grundler, F. M. (1997). Changes in the structure of *Arabidopsis thaliana* roots induced during development of males of the plant parasitic nematode *Heterodera schachtii*. European Journal of Plant Pathology, 103(2), 113–124. 10.1023/A:1008609409465

47. Stringlis, I. A., Proietti, S., Hickman, R., Van Verk, M. C., Zamioudis, C., & Pieterse, C. M. (2018). Root transcriptional dynamics induced by beneficial rhizobacteria and microbial immune elicitors reveal signatures of adaptation to mutualists. Plant Journal, 93(1), 166–180. 10.1111/tpj.13741

48. Sun, T., Zhang, Y., Li, Y., Zhang, Q., Ding, Y., & Zhang, Y. (2015). ChIP-seq reveals broad roles of SARD1 and CBP60g in regulating plant immunity. Nature Communications, 6(1), 10159. 10.1038/ncomms10159

49. Sun, Y., Fan, X. Y., Cao, D. M., Tang, W., He, K., Zhu, J. Y., He, J. X., Bai, M. Y., Zhu, S., Oh, E., Patil, S., Kim, T. W., Ji, H., Wong, W. H., Rhee, S. Y., & Wang, Z. Y. (2010). Integration of Brassinosteroid Signal Transduction with the Transcription Network for Plant Growth Regulation in *Arabidopsis*. Developmental Cell, 19(5), 765–777. 10.1016/j.devcel.2010.10.010

50. Taylor, N. G., Laurie, S., & Turner, S. R. (2000). Multiple cellulose synthase catalytic subunits are required for cellulose synthesis in *Arabidopsis*. Plant Cell, 12(12), 2529–2539. 10.1105/tpc.12.12.2529

51. Taylor-Teeples, M., Lin, L., De Lucas, M., Turco, G., Toal, T. W., Gaudinier, A., Young, N. F., Trabucco, G. M., Veling, M. T., Lamothe, R., Handakumbura, P. P., Xiong, G., Wang, C., Corwin, J., Tsoukalas, A., Zhang, L., Ware, D., Pauly, M., Kliebenstein, D. J., Dehesh, K., Tagkopoulos, I., Breton, G., Pruneda-Paz, J. L., Ahnert, S. E., Kay, S. A., Hazen, S. P., & Brady, S. M. (2015). An *Arabidopsis* gene regulatory network for secondary cell wall synthesis. Nature, 517 (7536), 571–575. 10.1038/nature14099

52. Urzúa Lehuedé, T., Berdion Gabarain, V., Ibeas, M. A., Salinas-Grenet, H., Achá-Escobar, R., Moyano, T. C., Ferrero, L., Núñez-Lillo, G., Pérez-Díaz, J., Perotti, M. F., Miguel, V. N., Spies, F. P., Rosas, M. A., Kawamura, A., Rodríguez-García, D. R., Kim, A. R., Nolan, T., Moreno, A. A., Sugimoto, K., Perrimon, N., Sanguinet, K. A., Meneses, C., Chan, R. L., Ariel, F., Alvarez, J. M., & Estevez, J. M. (2025). Two antagonistic gene regulatory networks drive *Arabidopsis* root hair growth at low temperature linked to a low-nutrient environment. New Phytologist, 245(6), 2645–2664. 10.1111/nph.20406

53. Völz, R., Kim, S. K., Mi, J., Mariappan, K. G., Guo, X., Bigeard, J., Alejandro, S., Pflieger, D., Rayapuram, N., Al-Babili, S., & Hirt, H. (2018). The Trihelix transcription factor GT2-like 1 (GTL1) promotes salicylic acid metabolism, and regulates bacterial-triggered immunity. PLoS Genetics, 14(10), 1–22. 10.1371/journal.pgen.1007708

54. Wu, T., Hu, E., Xu, S., Chen, M., Guo, P., Dai, Z., Feng, T., Zhou, L., Tang, W., Zhan, L., Fu, X., Liu, S., Bo, X., & Yu, G. (2021). clusterProfiler 4.0: A universal enrichment tool for interpreting omics data. The Innovation, 2(3), 100141. 10.1016/j.xinn.2021.100141

55. Wu, Z., Liang, S., Song, W., Lin, G., Wang, W., Zhang, H., Han, Z., & Chai, J. (2017). Functional and Structural Characterization of a Receptor-Like Kinase Involved in Germination and Cell Expansion in *Arabidopsis*. Frontiers in Plant Science, 8(November), 1–13. 10.3389/fpls.2017.01999

56. Wyss, U., & Zunke, U. (1986). Observation of the behaviour of second stage juveniles of *Heterodera schachtii* inside host roots. Revue de Nématologie, 9(2), 153–166.

57. Yang, J.-w., Kim, H. S., & Kim, Y.-H. (2026). Redox Regulation of Plant–Root-Knot Nematode Interactions: From ROS-Mediated Immunity to Sustainable Resistance. Antioxidants, 15(7), 853. 10.3390/antiox15070853

58. Yu, X., Li, L., Zola, J., Aluru, M., Ye, H., Foudree, A., Guo, H., Anderson, S., Aluru, S., Liu, P., Rodermel, S., & Yin, Y. (2011). A brassinosteroid transcriptional network revealed by genome-wide identification of BES1 target genes in *Arabidopsis thaliana*. The Plant Journal, 65(4), 634– 646. 10.1111/j.1365-313X.2010.04449.x

59. Zheng, B., Bai, Q., Li, C., Wang, L., Wei, Q., Ali, K., Li, W., Huang, S., Xu, H., Li, G., Ren, H., & Wu, G. (2022). Pan-brassinosteroid signaling revealed by functional analysis of NILR1 in land plants. New Phytologist, 235(4), 1455–1469. 10.1111/nph.18228

60. Zhu, A., Ibrahim, J. G., & Love, M. I. (2019). Heavy-Tailed prior distributions for sequence count data: Removing the noise and preserving large differences. Bioinformatics, 35(12), 2084–2092. 10.1093/bioinformatics/bty895

